# Investigating the impact of whole genome duplication on transposable element evolution in ray-finned fishes

**DOI:** 10.1101/2023.12.22.572151

**Authors:** Rittika Mallik, Dustin J. Wcisel, Thomas J. Near, Jeffrey A. Yoder, Alex Dornburg

## Abstract

Transposable elements (TEs) can make up more than 50% of any given vertebrate’s genome, with substantial variability in TE composition among lineages. TE variation is often linked to changes in gene regulation, genome size, and speciation. However, the role that genome duplication events have played in generating abrupt shifts in the composition of the mobilome over macroevolutionary timescales remains unclear. We investigated the degree to which the teleost genome duplication (TGD) shaped the diversification trajectory of the ray-finned fish mobilome. We integrate a new high coverage genome of *Polypterus bichir* with data from over 100 publicly available actinopterygian genomes to assess the macroevolutionary implications of genome duplication events on TE evolution. Our results provide no evidence for a substantial shift in mobilome composition following the TGD event. Instead, the diversity of the actinopterygian mobilome appears to have been shaped by a history of lineage specific shifts in composition that are not correlated with commonly evoked drivers of diversification such as body size, water column usage, or latitude. Collectively, these results provide a new perspective on the early diversification of the actinopterygian mobilome and suggest that historic ploidy events may not necessarily catalyze bursts of TE diversification and innovation.

## Introduction

Transposable elements (TEs) can account for over 50% of a vertebrate’s total genome content (Sotero-Caio et al. 2017). Aptly dubbed “jumping genes,” TEs possess the astonishing ability to rearrange and reposition themselves within a given genome (McClintock 1984), through the use of two primary strategies for transposing. Class I retrotransposons replicate through a “copy and paste mechanism” involving an RNA intermediate. This RNA molecule is reverse transcribed back into a DNA copy, which is then seamlessly integrated into the genome, leaving the original template element untouched (Hickman and Dyda 2015). In contrast, class II DNA transposons use a “cut-and-paste” technique to excise themselves from their current location and relocate to an alternate genomic location (Muñoz-López and García-Pérez 2010). Comparative studies of vertebrate TEs have revealed substantial heterogeneity in the composition of class I and class II TEs between major clades of vertebrates (Chalopin et al. 2015), These studies have implicated changes in TE composition with altered gene regulation (Torres et al. 2021; Rech et al. 2022), evolutionary changes in genome size (Böhne et al. 2008; Naville et al. 2019; Wong et al. 2019), evolutionary novelties (Senft and Macfarlan 2021), changes in life history (Niu et al. 2019), and speciation (Ricci et al. 2018; Serrato-Capuchina and Matute 2018) to name but a few. While the past decade has yielded tremendous strides towards developing an understanding of the general hallmarks of TE evolution, the evolutionary fate of TEs following whole genome duplication events is just beginning to emerge (Parisod et al. 2010; Rodriguez and Arkhipova 2018; Papon et al. 2023).

Genome duplication events have the potential to amplify genome complexity, with emerging evidence highlighting a possible role for duplication events enabling phenotypic evolution through duplicated genes (Moriyama and Koshiba-Takeuchi 2018). Studies within individual or closely related species that vary in their level of ploidy have revealed substantial modifications that can facilitate profound epigenetic repatterning within the domains of TEs. However, the impact of such changes on the mode of TE diversification remains unclear. For example, surplus gene copies can compensate for potential losses or modifications in expression caused by TE insertions, thereby facilitating extensive genomic modification through the actions of transposable elements. This redundancy hypothesis (Matzke and Matzke 1998) suggests a substantial shift in TE content and composition following a genome duplication event that continues as lineages diversify (Pariso d et al. 2010). An alternate hypothesis argues that genome duplication events correspond to a transitory phase for a species that is characterized by diminished population size (Lynch 2007). This bottleneck hypothesis argues that as the efficacy of selection decreases, the likelihood of moderately deleterious TE insertions within nascent polyploid genomes becoming fixed is increased. Consequently, this hypothesis would expect a pulse of TE diversification coincident with a genome duplication event (Parisod et al. 2010). Recently, a hypothesis framed around the concept of “Hopeful Monsters” suggested that the balance between the deleterious and beneficial aspects of TE proliferation can lead to occasional beneficial mutations that can promote speciation and contribute to the emergence of new traits (Turner et al. 2021). In contrast to the previous two hypotheses, this hypothesis predicts that genome duplications would result in only a limited number of TE changes, not a substantial pulse of diversification. Given that these hypotheses represent corner cases on a continuum of possibilities and alternate hypotheses, empirical comparative genomic studies are key to understanding the macroevolutionary implications of genome duplication events on the evolution of TEs.

Ray-finned fishes (Actinopterygii) offer an exemplar group for the macroevolutionary study of TEs and genome duplication events. Comprising half of all living vertebrate species, ray-finned fishes have successfully radiated across virtually all aquatic habitats including swamps (Albert et al. 2020), abyssal ocean depths (Davis et al. 2014; Miller et al. 2022), polar and high altitude regions (Near et al. 2012; Dornburg et al. 2017; De-Kayne et al. 2022; Parker et al. 2022) and caves (Wen et al. 2022). The evolutionary success of ray-finned fishes is unusual relative to other vertebrate groups. Over 99% of the over 35,000 living species of actinopterygians are teleosts, a clade that experienced a genome duplication event in its early evolutionary history (Brunet et al. 2006; Glasauer and Neuhauss 2014). In contrast, the three remaining extant ray-finned fish lineages – Holostei (gars and bowfin, 8 species), Acipenseriformes (sturgeon and paddlefish, 29 species), and Polypteridae (bichirs and ropefishes, 14 species)– did not undergo the teleost specific genome duplication event. Recent studies of holostean genomes have highlighted differences in the composition of TEs between holosteans and model teleosts such as zebrafish (*Danio rerio*) and medaka (*Oryzias latipes*) (Braasch et al. 2016; Thompson et al. 2021; Mallik et al. 2023), raising the possibility of an evolutionary shift following a genome duplication event in teleosts. Comparative genomic studies are needed to rule out the possibility that the mobilomes of holosteans, and not teleosts, are unusual relative to all other ray-finned fishes. To date, no comparative studies of the ray-finned fish mobilome have included a broad representative sampling of teleosts, holosteans, acipenseriforms, and polypterids. With the growing number of genomes of ray-finned fishes deposited in public data repositories, such studies are now possible.

Over twenty years ago, the *Takifugu rubripes* genome represented the first ray-finned fish genome sequenced (Aparicio et al. 2002). Decreased costs of sequencing have since led to a surge of efforts to sequence additional actinopterygian genomes (Fan et al. 2020; Randhawa and Pawar 2021) that now present a wealth of resources for phylogenetically comprehensive comparative studies. In addition to the availability of hundreds of teleost genomes, there are genomes of a few for non-teleost ray-finned fishes. For example, the sterlet (*Acipenser ruthenus*) genome revealed an independent historic genome duplication in this chondrostein lineage that provides an additional ancient duplication for understanding evolutionary patterns of TE diversification in actinopterygians (Cheng et al. 2019). Similarly, the sequencing of the Senegal bichir (*Polypterus senegalus*) leveraged the anatomy of polypterids to provide critical insights into how vertebrates achieved the transition from water to land (Bi et al. 2021). In addition, the recently sequenced genome of a second polypterid, *Erpetoichthys calabaricus* [Reedfish; Assembly (ErpCal1.1; NCBI Annotation Release 100)], has been deployed alongside the *P. senegalus* genome to provide further insights into other aspects of early vertebrate diversification that include the evolution of keratins (Kimura and Nikaido 2021), olfactory receptors (Zhang et al. 2021), and numerous other traits (Helfenrath et al. 2021; Mikami et al. 2022). However, based on BUSCO scores, the currently sequenced genome of *P. senegalus*, remains incomplete. As the only *Polypterus* genome sequenced, the lack of an additional *Polypterus* genome challenges interpretation of results concerning the distribution of TEs in this lineage. Given the phylogenetic position of polypterids, additional genome sequencing efforts are of extreme value for contextualizing the diversification of ray-finned fish TEs.

Here we present a high-quality chromosome-level assembly and annotation for an additional *Polypterus* species, *Polypterus bichir*. We integrate analyses of TEs within this genome with data on TE content for all other major lineages of actinopterygians to investigate the impact of genome duplication on the early evolution of the teleost mobilome. Using a comparative phylogenetic framework, we test for associations between genome size and the composition of the mobilome, variations in mobilome composition between teleosts and non-teleost ray-finned fish lineages, as well as possible correlations between TE content and aspects of actinopterygian biodiversity such as habitat, body size, or water column occupation. We additionally reconstruct the ancestral mobilome of actinopterygians through the TGD event and assess the phylogenetic signal of Class I and Class II TEs. These results provide a new perspective on the early diversification of the actinopterygian mobilome and the impact of the TGD on its evolution.

## Results and Discussion

### Bichir genomes: an example of evolutionary conservation or recent divergence?

We present a high-quality assembly of the *Polypterus bichir* genome (NCBI Bioproject PRJNA811142). The results of a BLAST search of mitochondrial COI from our specimen against polypterid barcode sequences on NCBI verified the identification as *Polypterus bichir lapradei*, a currently not recognized subspecies based on morphology that has been suggested to represent a genetically distinct lineage (Near et al. 2014). The 10X supernova assembly from Dovetail Genomics (Scotts Valley, CA) resulted in 130,773 scaffolds forming a total final genome size of 3,962,089,718bp. 13.9% of the genome (550,533,170bp) is composed of the ambiguous base “N” and a GC content of 39.21%. During Dovetail Hi-Rise assembly, the input assembly was further incorporated into 70,587 longer scaffolds. The total length of the resulting Dovetail Hi-Rise assembly was 3905.43 Mbp, with a contig N_50_ of 37.39 kbp. The N_50_ of the assembly was 202.693 Mbp scaffolds with a L_50_ of seven scaffolds. This is similar to the *E. calabaricus* genome which is 3.6Gb long, with a contig N_50_ size of 6.8 Mb kbp, and a scaffold N_50_ size of 217.7 Mbp, and the *P. senegalus* reference genome, which is 3.7 Gbp, with a scaffold N_50_ of 189.69 Mbp, and contig N_50_ of 4528.14 kbp (**Supplemental Table S1**). However, a comparison of the BUSCO analysis on the *P. senegalus* reference genome to the newly sequenced *P. bichir* genome reveal a striking difference between the two assemblies (**Supplemental Table S2** and **Supplemental Figure S1**). We find nearly triple the number of actinopterygian orthologs in the *P. bichir* genome relative to the *P. senegalus* genome (**Supplemental Figure S1**). This suggests that the *P. bichir* sequence may fill additional gaps in our understanding of the genomic evolution of polypteriforms outside of the mobilome elements discussed in this study.

Contrasting patterns of synteny between our new sequenced genome and those of *P. senegalus* and *E. calabaricus* using D-genies supports a high level of synteny between these species (**Supplemental Figure S2**). Further, the number of *P. bichir* superscaffolds (18) reflect the described karyotypes of most polypterids: *E. calabaricus* (n=18)*, Polypterus palmas* (n=18)*, Polypterus delhezi* (n=18), *P. senegalus* (n=18), and *Polypterus ornatipinnis* (n=18) (Bachmann et al. 1972; Vervoort 1980; Morescalchi et al. 2007; Krysanov and Golubtsov 2014). The only exception to this chromosome count known in polypterids is in *P. weeksi* (n=19) (Vervoort 1980), the sister lineage to *Polypterus ornatipinnis*. While closely related vertebrate taxa can often diverge substantially in their TE content over time, we find that this is not the case in bichirs. Instead, the conservation of karyotype and synteny across these species is also reflected in the relative abundances of TEs across these species. For *P. bichir*, RepeatModeler quantified 53.40% of the genome to be composed of transposable elements. This is similar to *P. senegalus* with (54.66%) and *Erpetoichthys calabaricus* (60.35%) and overall posits a surprising level of genomic conservation between species of polypterids. Such conservation could be a consequence of multiple factors such as slow rates of molecular evolution such as those observed in gars and Bowfin (Wright et al. 2022), the possible geologically recent crown age of Polypteridae estimated using molecular clocks (Near et al. 2014), or a combination of the two to name but one set of possibilities. Regardless of the mechanism, our sequencing of the *P. bichir* genome reveals polypterids as a candidate lineage for future studies of the mechanisms that promote genomic stability within a clade.

### The evolution of the ray-finned fish mobilome

Placing our characterization of the polypterid mobilome into the context of other major ray-finned fish lineages reveals a complex history of mobilome evolution over the past 400 million years (**Figure 1A**). The reconstructed phylogenetic history of TE evolution in actinopterygians is not consistent with a sudden TE proliferation coincident with the TGD event. Instead, our analyses reveal that compositional TE patterns at this scale largely reflect shifts within actinopterygian subclades. For example, cichlids have similar compositional patterns relative to pufferfish, the latter of which exhibits an increase in the number of LINE elements (**Figure 1A**). Independently, Lampriformes (*Regalecus* & *Lampris*) also exhibit an expansion of their LINE elements to a relative level similar to that observed in pufferfishes (**Figure 1A**). The ubiquity of such patterns of clade-specific heterogeneity are strongly supported by tests of phylogenetic signal using both Pagel’s lambda (λ) and Blomberg’s K (K) (**Supplemental Table S3**). In all cases, K values are significant (LINE | *K=0.945, p=1e-04*; DNA | *K=0.478, p=6e-04*; SINE | *K=0.529, p=3e-04*; and LTR | *K=0.405, p=1e-04*). Likewise, quantifications of lambda values for DNA (*λ=0.98, p=1.89e-13*), LTR (*λ=0.819, p=6.64e-11*), LINE (*λ=0.975, p=1.51e-19*), and SINE (λ∼1.0, p=4.37e-15) are very close to 1, indicating strong phylogenetic signal, supporting a pattern of similar relative TE abundances between closely related taxa and increasing disparity between subclades.

**Figure 1.**
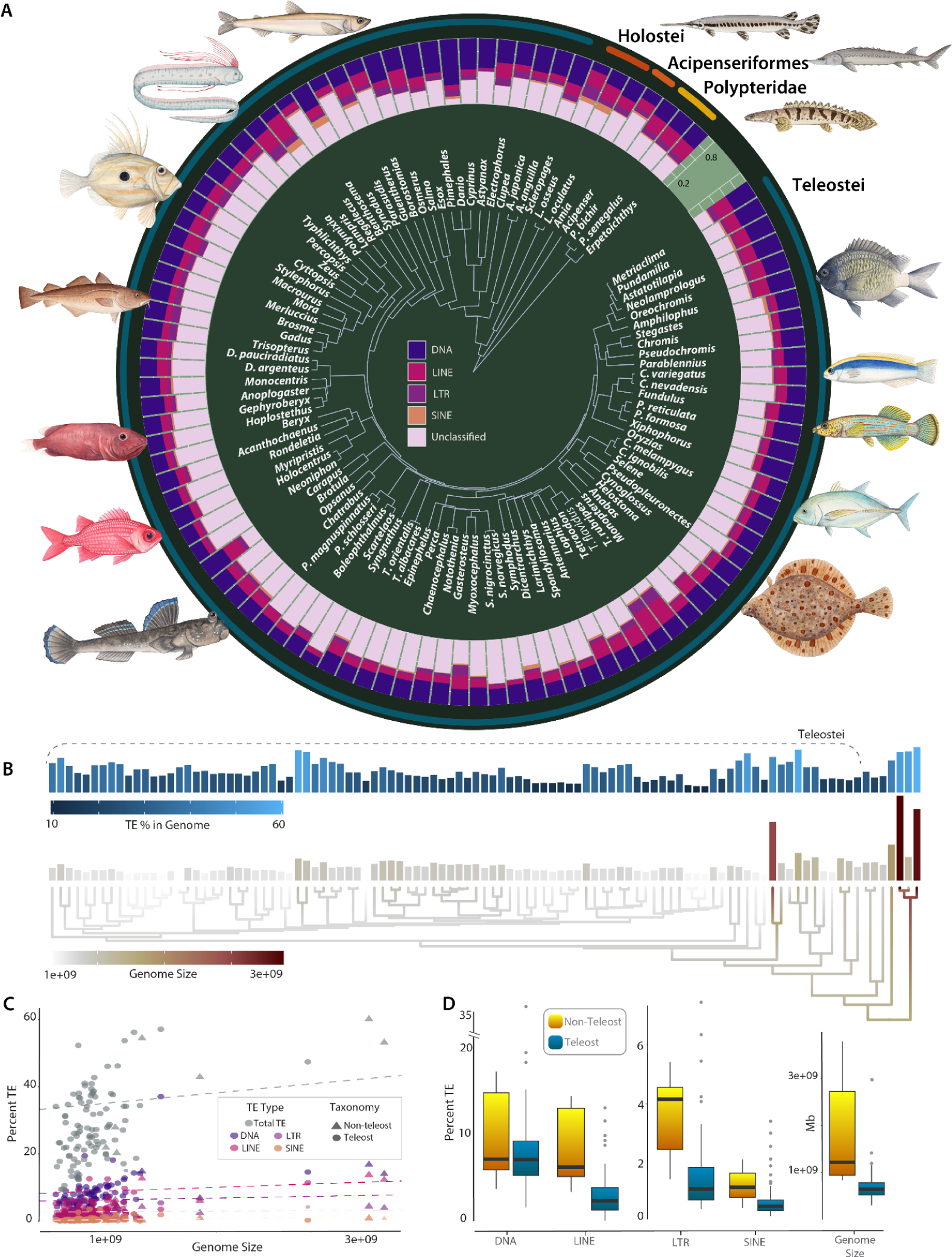
Major patterns of TE evolution across the evolutionary history of ray-finned fishes. (A) The relative abundance of DNA, LTR, LINE, SINE transposons, and unknown transposons relative to actinopteryigiian phylogeny are shown. (B) Total TE % and absolute genome size in the context of evolutionary history are compared. (C) Results of a phylogenetic regression to assess the relationship between genome size and TE abundance are provided. (D) TE % and genome size between teleosts and non-teleost actinopterygians are compared. Colors in A correspond to elements labeled in the panel. Colors in B are shaded relative to high (light blue or red) and low values (dark blue, or beige) in the upper and lower panels respectively. Colors in C correspond to the labels in A for individual elements. Panels in D correspond to the quantiles of each category with dark horizontal lines indicating the mean value.

Contrasting relative TE content (**Figure 1B**) against total genome size reveals numerous expansions and contractions of both genome size and relative TE abundances across the phylogenetic diversity of ray-finned fishes. Importantly, there is no signal of a shift in the rate of TE evolution with the origin of teleosts. Instead, the reconstructed history indicates that expansions and contractions of both relative TE abundance and genome size appear to heterogeneously occur across various teleost and non-teleost lineages. Phylogenetic regression analyses indicate that shifts in genome size are weakly correlated with changes in the relative abundances of TEs (**Figure 1C; Supplemental Table S3**). Correspondingly, we find strong evidence that genome size in ray-finned fishes also exhibits a pattern of strong phylogenetic signal (*K=0.659, p=6e-04; λ=0.717, p=4.45e-15*), paralleling a major trend in genome size evolution that has been found across the Tree of Life. Lineages as disparate as angiosperms (Beaulieu et al. 2008; Alonso et al. 2015), *Drosophila (Sessegolo et al. 2016)*, marine dinoflagellates (Lo et al. 2022), bacteria and Archaea (Martinez-Gutierrez and Aylward 2022) exhibit strong phylogenetic signal in genome size evolution. Given that the relative abundance of TE content in a genome is weakly correlated with genome size in ray-finned fishes (Lehmann et al. 2021)**Figure 1D**; and numerous other lineages (Chalopin et al. 2015), it is possible that TEs evolution exhibits a signature of strong phylogenetic signal across the Tree of Life (Dodsworth et al. 2015).

Multiple factors promote genome size evolution (Olmo et al. 1989), including hypothesized correlations between the molecular evolution of the mobilome and overall increases in genome size. For example, transposition rates generally exceed excision rates (Kelleher et al. 2020), enabling TEs to contribute to the enlargement of genomes and exert an influence on genome size evolution. However, whether the TGD event resulted in a shift in the mode of evolution between the teleosts and non teleosts mobilome remains unclear. Using a phylogenetic analysis of variance (ANOVA), we find no support for a significant difference in TE content between teleosts and non-teleosts (p = 0.06; **Figure 1D**). Additionally, phylogenetic regression analyses strongly support a correlation between the relative content of all mobilome elements, indicating that the increase or decrease in one will also result in an increase in the other (**Figure 2**). These results suggest that the overall relative composition of the ray-finned fish mobilome has been shaped more by recent lineage specific shifts in genome evolution than historical legacy stemming from the TGD event.

**Figure 2.**
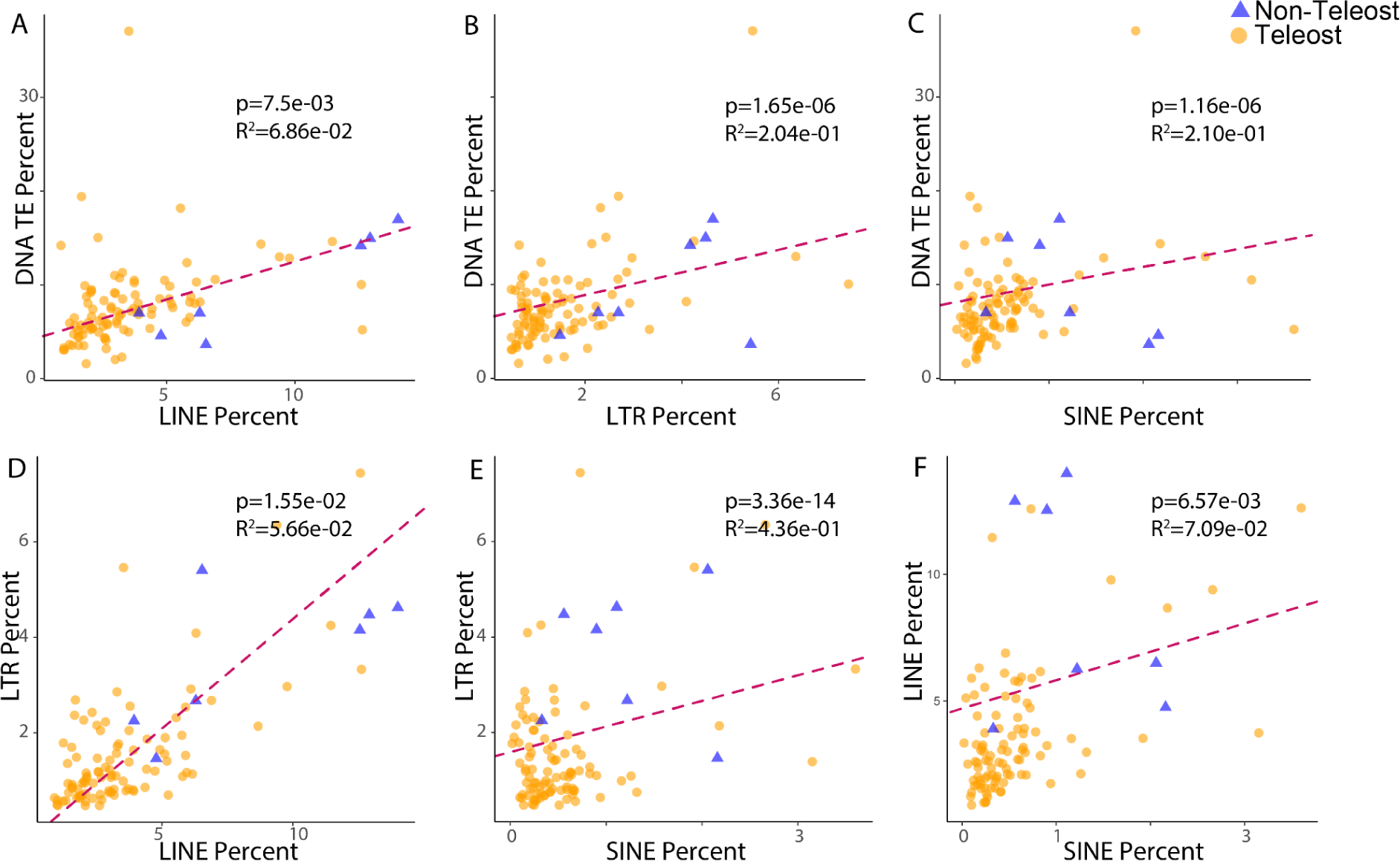
Correlation between relative class I and class II mobilome content. Panels (A), (B), and (C) display the correlations between DNA transposons and LINE, LTR, and SINE transposons, respectively. Panels (D) and (E) illustrate the correlations between LTR transposons and LINE and SINE transposons, respectively. Panel (F) presents the correlation between LINE and SINE transposons.

### Concerning the TGD and the appearance of novel mobilome elements

It is possible that the TGD catalyzed the evolution of major groups of novel TE’s. However, when placing the twenty-five main superfamilies of the teleost mobilome (Chalopin et al. 2015) into the context of the phylogenetic diversity of ray-finned fishes, we reveal that this effect was muted. We estimated the ancestral mobilome of actinopterygians utilizing a model-averaged stochastic character mapping approach (Revell 2023). Out of 25 superfamilies, the presence of 24 superfamilies observed in teleosts is mirrored within non-teleosts (**Figure 3**). The only exception to this is the DNA transposon superfamily Chaepev (Kapitonov and Jurka 2007). It is unlikely that this superfamily arose as a consequence of the TGD, as it is present in several arthropods as well as a range of vertebrate species including anoles (Novick et al. 2011), and lampreys (Zhang et al. 2014). Chapaev sequences have also been documented from White Sturgeon (*Acipenser transmontanus*) (Zhang et al. 2014), suggesting that additional species of non-teleost ray-finned fishes possess this superfamily.

**Figure 3.**
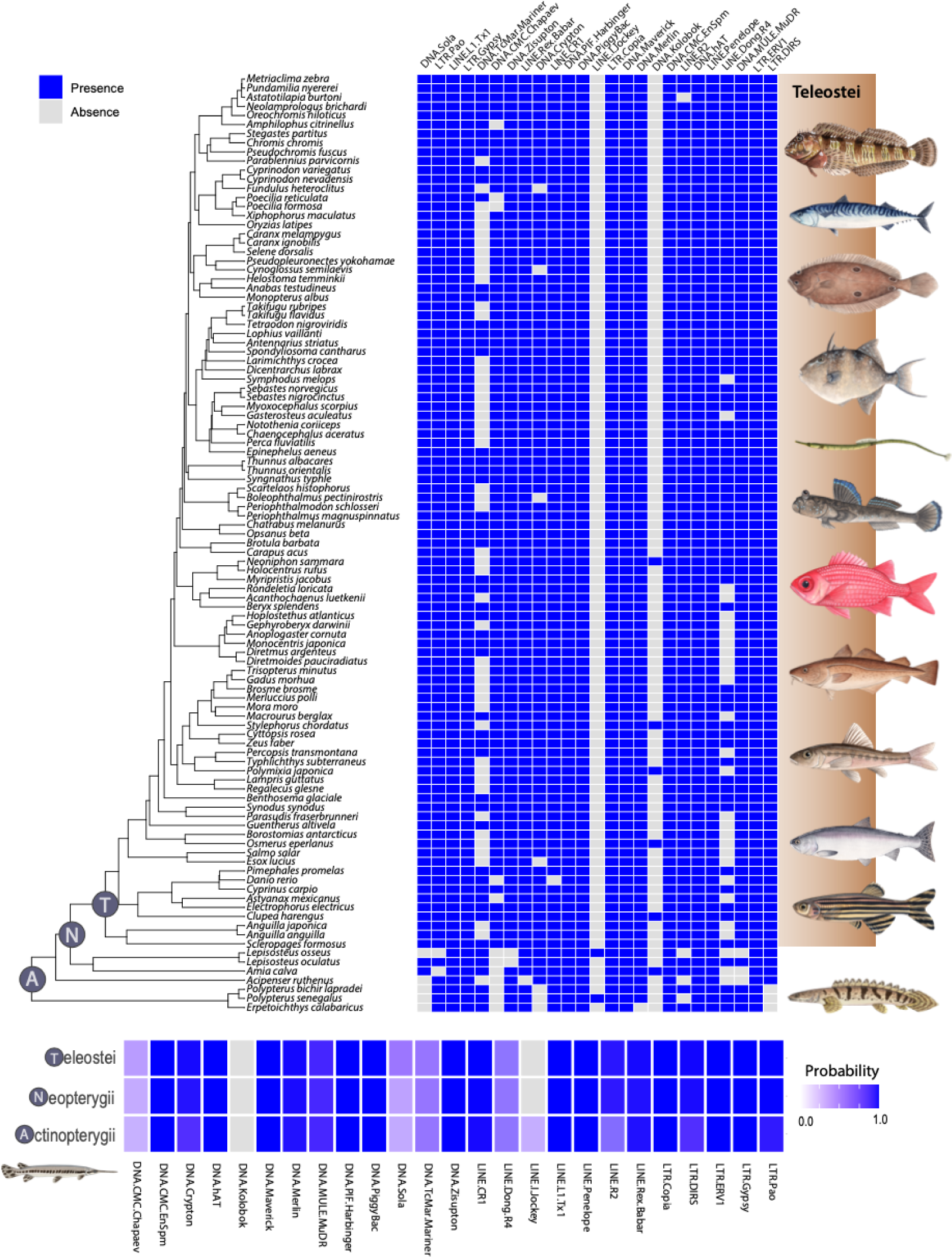
Reconstructing the ancestral Actinopterygian mobilome. The presence and absence of transposable element (TE) superfamilies (listed above) are represented by blue and grey squares, respectively, for various species on the left (top panel). The brown-shaded area represents teleosts, while the unshaded region below represents non-teleosts with evolutionary relationships between taxa depicted by the phylogenetic tree on the far-left. This input data was used to estimate the ancestral mobilome (the probability of each TE superfamily being present or absent) for the most recent common ancestor of all Actinopterygii (A), Neopterygii (N), or Teleostei (T) using SIMMAP (bottom panel) With a resulting heatmap plotted using ggplot (Wickham 2016).

Genome duplication events are hypothesized to form a substrate for genomic innovation (Crow et al. 2006). In contrast to this expectation, our results reveal numerous instances of numerous likely TE losses as well as independent gains. For example, we find repeated losses of TE super families such as LINE Dong, DNA TcMariner, and DNA Crypton in teleosts (**Figure 3**). LTR-Pao is absent in Longnose Gar and Bowfin, suggesting a loss in holosteans. LTR-DIRS are absent in all the three polypterids (*P. bichir, P. senegalus* and *E. calabaricus*). In addition, we identify the TE superfamily Jockey, a LINE element, exclusively in *Lepisosteus osseus* and *P. senegalus*. Jockey has been previously confirmed in coelacanth (Chalopin et al. 2014) and lampreys, and this marks the first reported instance of its presence in actinopterygians. It is possible that this element has been lost multiple times independently, however Jockey is known to have high rates of horizontal transfer in other lineages (Reiss et al. 2019; Tambones et al. 2019), raising the possibility that these were independent gains.

Studies of vertebrate genome evolution often assume vertical transmission as the dominant mode of evolution. However, recent work has highlighted the impact of horizontal transfer events in the evolution of the vertebrate mobilome (Zhang et al. 2020; Galbraith et al. 2022; Paulat et al. 2023). Our ancestral mobilome reconstructions of the DNA Kolobok superfamily are in line with phylogenetic patterns expected by a model of horizontal transfer (**Figure 3**). In our sampling, the presence of Kolobok is limited to five distantly related teleost lineages, as well as the Longnose Gar (*Lepisosteus osseus*) among non-teleost ray-finned fishes. It is possible that the Kolobok element has been repeatedly transferred throughout the evolutionary history of Actinopterygii. Such transfer events are considered more likely by recent observations of horizontal transfer at more recent time scales. For example, there is evidence of horizontal gene transfer of DNA transposons like Merlin, TcMariner, and PiggyBAC in salmonids (de Boer et al. 2007). Likewise, there has been evidence of horizontal transfer of Chapaev transposons in White Sturgeon, Pacific Bluefin Tuna (*Thunnus orientalis*) and Blue Catfish (*Ictalurus furcatus*) (Zhang et al. 2014). As the number of high quality genomes for species of ray finned fishes continue to accumulate, future studies of the extent of horizontal transfer events across the ray-finned fish mobilome offer an exciting research prospect.

### Lineage-specific expansions of the teleost mobilome

Transposable elements (TEs) likely play a beneficial role by enhancing an organism’s ability to respond to dynamic environmental conditions (Yuan et al., 2018). Numerous studies have linked TE activity to an organism’s responsiveness to environmental factors (Capy et al. 2000; Fujino et al. 2011; Hua-Van et al. 2011; Baduel et al. 2021). This association between TEs and environment suggests that there may be a correlation between relative TE abundance and specific aspects of an organism’s ecological niche or life history. However, we find no such association for several factors often invoked to explain diversification patterns at this evolutionary scale. Under the Brownian motion model, our phylogenetics least squares regression (PGLS) analysis between TE content and maximum body size revealed no evidence for a strong correlation between these traits (**Figure 4A** and **Supplemental Tables S4, S5, S6 and S7**; Likewise, we find no statistically supported correlation between TE content and depth occupancy in marine fishes (**Figure 4B**). There is also no general pattern of a correlation between TE content and the average latitude of a species geographic distribution (**Figure 4C**). The only exception to this lack of correlation between latitude and TE content occurs in SINEs (**Supplemental Table S6**) under the Brownian motion model (p=0.0197). This correlation is likely driven by a reduction in SINE elements in Antarctic notothenioids, which experienced an unusual bout of genomic evolution prior to their diversification in the Southern Ocean (Daane et al. 2019). Regardless, depth, body size, or latitude appear not to be predictors of mobilome evolution, even when differentiating between marine, freshwater, or estuarine fishes at this evolutionary scale. In all cases, phylogenetic regression results were largely similar between a brownian and Orstein-Uhlenbeck model of trait evolution (**Supplemental Table S6 & S7**).

**Figure 4:**
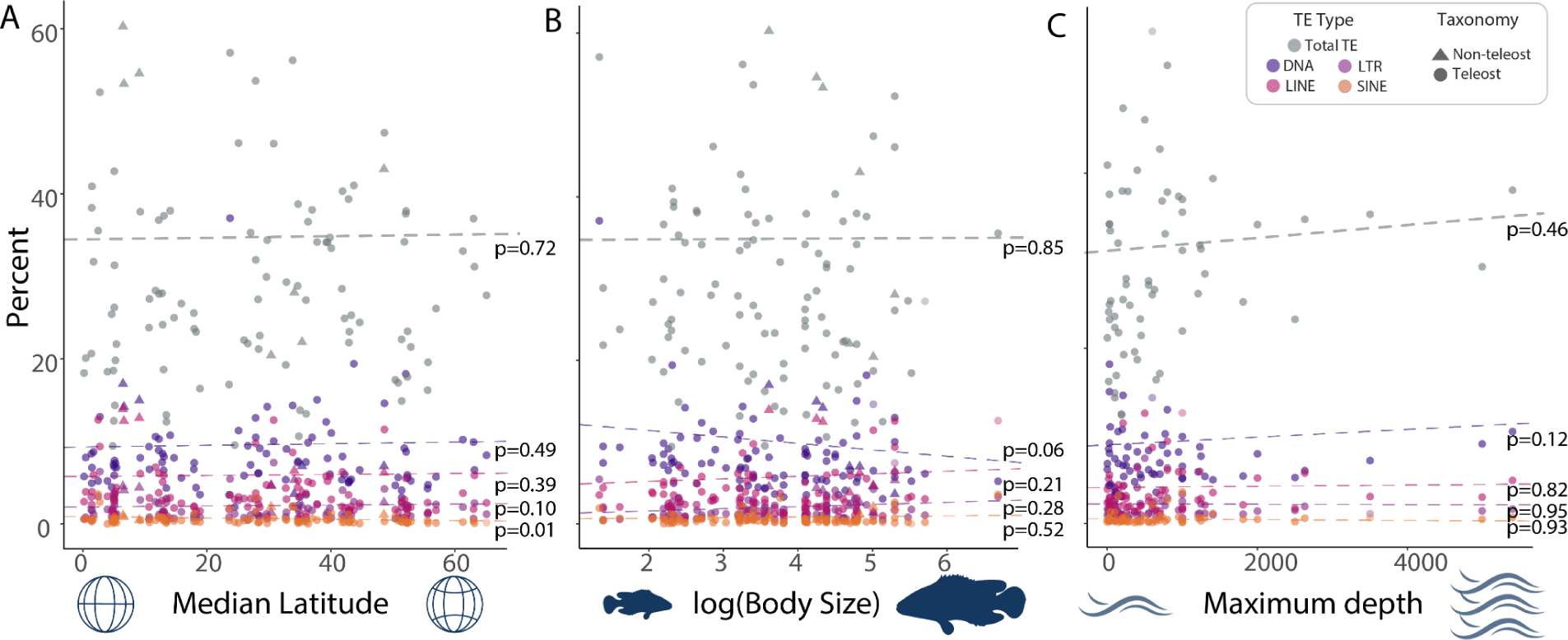
Correlation of TEs with abiotic factors. (A) This panel displays the results of phylogenetic generalized least-squares (PGLS) regressions between TE percentages and median latitude (B) This panel portrays the PGLS regression results between TE percentage and body size (log-transformed). Lastly, (C) depicts PGLS results between TE abundance and the maximum habitat depth of the species. The colors and shapes of each data point on the plot are defined in the key.

The TGD certainly could have presented opportunities for a burst of genomic novelty. However, the next 300 million years of ray-finned fish evolution would certainly be expected to shape the genomes of different lineages responding to different biotic and abiotic conditions. As such it is possible that the signal of the genome duplication has either eroded, or that rapid rates of mobilome evolution manifest as high between-clade heterogeneity at the scale of all ray-finned fishes. Clade-specific changes in the mobilome are readily apparent when considering a phylogenetic principal components analysis (PCA) of mobilome abundances (**Figure 5**). For example, PC1 corresponds to sharp divergence between *Esox*, salmonids, and three bichirs, from acanthomorphs (**Figure 5 and Supplementary Figures S4, S5, S6**). This alignment of taxa with PC1 values coincides with the prevalence of high percentage of DNA transposons that comprise the transposable elements within fish genomes that influence genome size variation among teleosts such as zebrafish (**Supplemental Figure S3)**, medaka, stickleback, and *Tetraodon* (Jiang et al. 2016). It is certainly not possible to discount a possible role for life history shifts shaping the mobilome among closely related species or that differences between Class 1 and Class II replications serve as an opportunity for substantial expansion of the mobilome.

**Figure 5:**
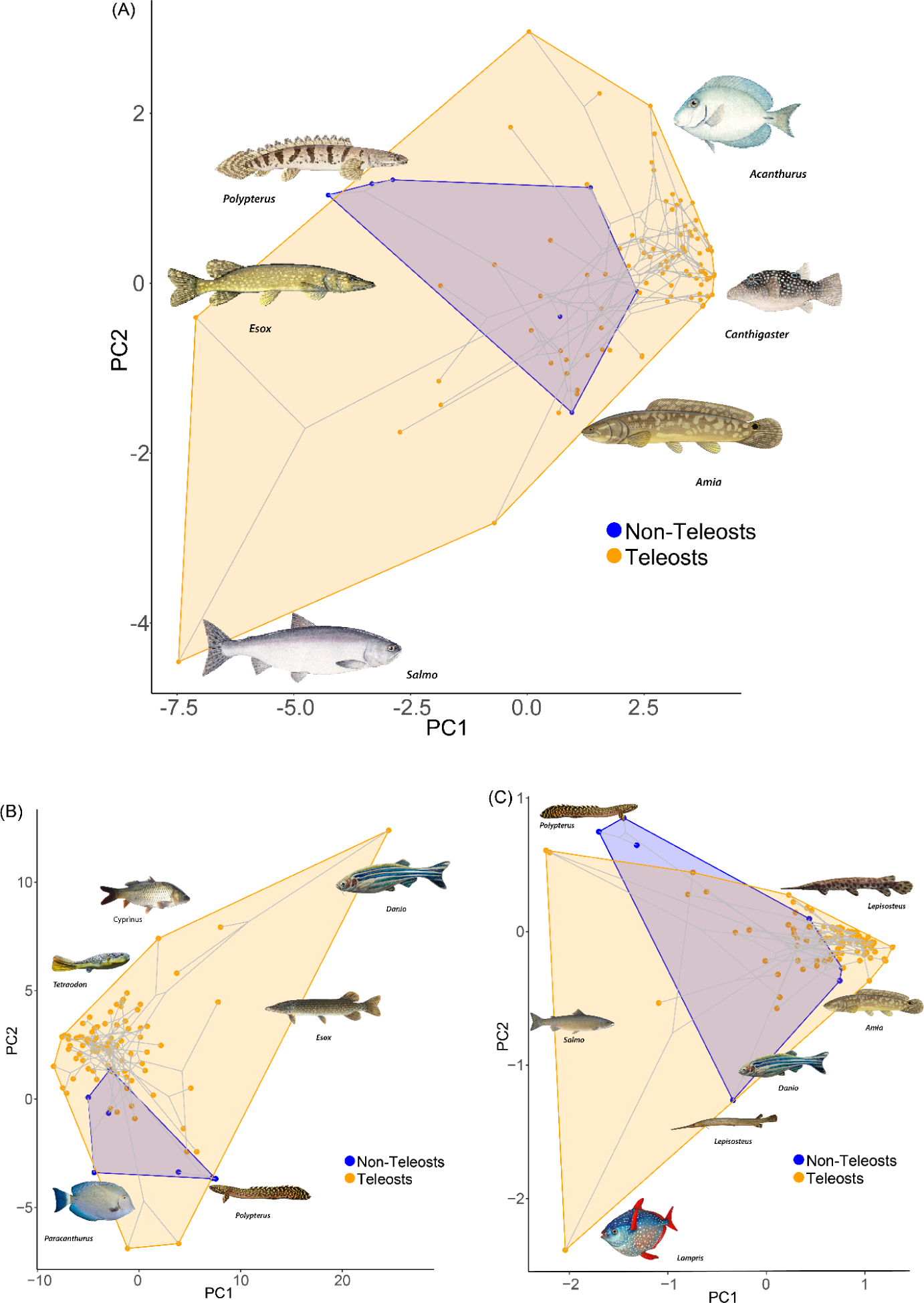
Considering Transposable Element (TE) diversity in the context of evolutionary history. (A) Results of a Phylogenetic PCA on the abundances of LTRs, DNA transposons, LINEs, and SINEs that project the phylogeny and points onto a 2D plot of PC1 and PC2. (B) depicts the resulting space based on the residual variation following linear regression between major TE classes and genome size. (C) Depicts the results of a PCA on the residual variation of the abundances of all 25 superfamilies accounting for differences in genome size, consisting of 13 DNA transposons, 7 LINEs, and 5 LTR subfamilies. Blue indicates non-teleost actinopterygians, orange indicates teleosts. All the images of the fishes are from Wikimedia commons.

### Conclusion

Our results shed light on the potential impact of the TGD on the diversification of the ray-finned fish mobilome. We find no evidence for a discernible temporal signature of diversification coincident with the TGD, raising the possibility that this genome duplication event did not exert a long-lasting influence on transposable element diversification. The absence of a discernible temporal signature of diversification coincident with the TGD raises important questions about the role of ploidy in TE diversification. Our results suggest that the ray-finned fish mobilome appears to maintain a consistent profile, with substantial similarity between the mobilomes of teleosts and non-teleost ray-finned fishes. However, mobilome content between taxa varied substantially, from around 55% in Zebrafish to about 6% in pufferfish genomes. These shifts cannot be entirely explained by differences in genome size as genome compaction in smooth pufferfish like *T. rubripes* and *Tetraodon nigroviridis* is not associated with an overall reduction of retrotransposon diversity (Volff et al. 2003). Similarly, Medaka and Zebrafish have been found to have very similar L1 retrotransposon diversity, despite a large disparity in genome size (Kordis et al. 2006). Our results strongly imply that the history of the ray-finned fish mobilome is one of lineage-specific evolution, suggesting future studies between more closely related clades to be particularly fruitful as genomic resources become available. These results also place ray-finned fishes into the overall history of the vertebrate mobilome.

Early diverging aquatic vertebrates (e.g., coelacanth, ray finned fishes, cartilaginous fish, and lamprey) contain a significantly broader spectrum of mobilome diversity in the genome relative to birds or mammals, ranging from 22 to 27 TE superfamilies. Among these, certain autonomous elements like ERVs, LINE1 retrotransposons, TcMariner, or hAT DNA transposons, as well as non-autonomous ones like V-SINE (Piskurek and Jackson 2011), are widely distributed across all the vertebrates. This suggests their likely presence in ancestral vertebrate genomes. Conversely, some TE superfamilies have likely been lost or are on the path to extinction in specific lineages, L2 and Helitrons in birds (Chalopin et al. 2015) and gypsy retrotransposons in birds and mammals (Volff et al. 2003) fall into this category. Further diversity can be compared between lineages with respect to TE superfamilies. The LINE1 retrotransposon family comprises 20% of human and mouse genomes. Mammalian species have LINE1 transposons that have a large number of shared sequences, making it difficult to distinguish the different types of L1 (Ivancevic et al. 2016), and hence are grouped together as RT-family. In the case of ray-finned fishes, there is evidence of both diverse L1/Tx1 actinopterygian specific families as well as the presence of mammalian L1 (Sotero-Caio et al. 2017). In contrast, we see a lower distribution of endoretroviruses (ERVs) in actinopterygians than mammals and birds, with the amount ranging from 0.033% in Fugu to 0.76% in zebrafish (Carducci et al. 2020). These contrasts parallel our findings of clade-specific heterogeneity within ray-finned fishes, and underscore the potential for other factors commonly invoked in biodiversity studies (i.e., extinction events) to have shaped mobilome divergences between ray-finned fish lineages and other vertebrates. It is clear that we are only beginning to unmask the complexity of transposon dynamics and decipher the intricate processes that contribute to TE diversification over deep evolutionary timescales.

## Methods and Materials

### Sample acquisition and sequencing

All research involving live animals was performed in accordance with relevant institutional and national guidelines and regulations, and was approved by the North Carolina State University Institutional Animal Care and Use Committee (IACUC). We acquired an individual *Polypterus bichir* specimen (Yoder lab ID 0051) through the pet trade that was anesthetized using MS-222 for the extraction of blood (10 ml). The fish was then euthanized for dissection of tissue samples and the voucher specimen was deposited in the North Carolina Museum of Natural Sciences Ichthyology Collection (NCSM 111902). Blood was shipped to Dovetail Genomics, LCC (Scotts Valley, CA) for genomic DNA extraction, library preparation, sequencing, and assembly. Samples were extracted by Dovetail staff using Qiagen and Cell Culture Midi Kit (Qiagen, Gmbh), yielding DNA with an average fragment length of 95 kbp that was used in the construction of HiC sequencing libraries.

### Chicago Library preparation and sequencing

A Chicago library was prepared using the methods as described in (Putnam et al. 2016). About 500 ng of High Molecular Weight genomic DNA, with mean fragment length of 95 kbp. Chromatin was reconstituted in vitro by incorporating the DNA with purified histones and chromatin assembly factors and then fixed by formaldehyde. *Dpn*II was used to digest the chromatin, followed by filling in the 5′ overhangs with biotinylated nucleotides, and then ligating the free blunt ends. After the ligations step, DNA was purified from protein by reversing the crosslinks. Biotinylated free ends were removed from the purified DNA by treating it. The DNA was sheared to ∼350 bp fragment size and Illumina compatible adapters along with NERNext Ultra enzymes were used to generate sequencing libraries. Streptavidin beads were used ahead of PCR enrichment of each library to isolate the biotin-containing fragments. The libraries were sequenced on an Illumina HiSeqX providing 30.58x physical coverage of the genome (1-100 kbp).

### Dovetail Hi-C library preparation and sequencing

The preparation of a dovetail Hi-C library was executed as described (Lieberman-Aiden et al. 2009). For each library, the chromatin was fixed in place in the nucleus by crosslinking with formaldehyde, and the extracted fixed chromatin was then digested with the restriction enzyme *Dpn*II that produces 5′ overhangs. These 5′ overhangs were filled in with biotinylated nucleotides, and then the resulting blunt ends were ligated. Crosslinks were then reversed to obtain the purified DNA, and which was treated to remove excess biotin. A Hi-C library was then created by shearing the DNA to ∼350 bp mean fragment size. Sequencing libraries were generated using NEBNext Ultra enzymes and Illumina compatible adapters. Streptavidin beads were used prior to PCR enrichment of each library to isolate the biotin containing fragments. Libraries were sequenced using an Illumina HiSeqX which yielded 191 million paired end reads (2×150 bp) and provided 8,427.66 x physical coverage of the genome (10-10,000 kbp).

### Scaffolding the assembly with Hi-Rise

Sequencing reads were assembled with Hi-Rise, a software pipeline designed to scaffold genome assemblies using proximity ligation data (Putnam et al. 2016). It uses the *de novo* assembly, shotgun reads, Chicago library reads and Dovetail HiC library reads as input and conducts an iterative analysis. The shotgun and Chicago library sequences are first aligned to the draft input assembly using a modified SNAP read mapper (http://snap.cs.berkeley.edu), with modifications such as masking out the base pairs that followed a restriction enzyme junction. Hi-Rise then modeled the separations of Chicago read pairs mapped within draft scaffolds, using a likelihood model for genomic distance between read pairs. The likelihood model produced by HiRise was used to identify and break putative misjoins and also score and make prospective joins. Dovetail HiC library sequences were analyzed using the same methods after aligning and scaffolding the Chicago data. Once all the sequences were aligned and scaffolded, shotgun sequences were used to close the gaps between contigs.

### Contamination removal and species verification

Dovetail staff have noted that when pooled with other samples on Illumina sequencing platforms, 10X Chromium Genome solution libraries are susceptible to a small degree of index hopping that can result in minor incorrect assignment of sequenced reads during demultiplexing. This low level of index misassignment, typically results in sequence contaminants impacting, but limited to, small scaffolds (typically less than 10 kb) in the final assembly. To mitigate this, dovetail staff leveraged any uncharacteristic number of reads per barcode associated with impacted scaffolds to identify and reliably isolate them from the final assembly. This was accomplished by aligning the 10X reads to the supernova assembly, recording the barcode count for the aligned reads, and recording the number of reads that aligned to each scaffold and were tagged with the same barcode. The median number of reads per barcode for each scaffold were then calculated and scaffolds with a distinct anomalously low ratio were removed.

Currently *Polypterus bichir* is described as a single species with the subspecies *Polypterus bichir bichir* and *Polypterus bichir lapradei* no longer considered valid (Moritz and Britz). This sinking of subspecies is a consequence of the high degree of morphological similarity (Britz and David Johnson 2003; Britz and Johnson 2010). However, molecular investigations have suggested the possibility that *P. bichir bichir* and *P. bichir lapradei* may be genetically distinct (Suzuki et al. 2010) and these have been treated as independent lineages (Near et al. 2014). As no genetic species delimitation analyses have been conducted between these putative genetic lineages, we extracted the mitochondrial barcode COI from our assembly and used a BLAST search against polypterid sequences on NCBI to verify identification that included barcodes from *Polypterus bichir bichir* and *Polypterus bichir lapradei*. This ensured that we accounted for possible future taxonomic revisions while remaining consistent with current taxonomy.

### RNA sequencing and assembly

RNA was extracted (Qiagen RNeasy kit) from the spleen, kidney, gill, heart, eyes and intestine of the same individual *P. bichir* as the genome sequence. Quantity and integrity of the isolated RNA were assessed using a NanoDrop 1000 (Thermo Fisher) and Agilent Bioanalyzer respectively. The process of mRNA enrichment was done using oligo(dT) beads, and rRNA was removed using a Ribo-Zero kit (Epicentre, Madison, WI). Each RNA extraction was equalized for a final concentration of 180 ng/µL. Library preparation and sequencing was performed by Novogen Corporation (Sacramento, CA). Next-gen sequencing (2 × 150 bp paired end reads) was performed on a NovaSeq 6000 instrument (Illumina). Adapter sequences and poor quality reads were filtered with Trimmomatic v34 (Bolger et al. 2014). The transcriptome was de novo assembled with Trinity v2.11.0 (Grabherr et al. 2011) followed by BUSCO analysis to assess completeness of the transcriptomes (Manni et al. 2021). Raw reads and computationally assembled transcriptome sequences were deposited onto NCBI under the accession numbers SRR19537224 - SRR19537230 and GKOV01000000 respectively.

### Gene ontology assessment

We conducted a series of analyses for gene ontology assignment. First, the RNA-seq data assembled using Trinity was translated with Transdecoder, enabling the identification of potential coding regions in the transcript sequences. The longest open reading frames (ORFs) were extracted and subjected to BLASTx and BLASTp analyses against the Uniprot database (November 2021 release), yielding the top target sequences for each transcript. Subsequently, Hmmscan v.3.3.2 was employed to search for protein sequences in the Pfam-A database (November 2021 release). Signalp v.5.0b and TMHMM v.2.0c were used to detect signal peptides and transmembrane proteins, respectively. Trinotate v.3.2.2, in combination with the Trinity assembled transcriptome and the longest ORFs from Transdecoder, generated a gene transmap. This process facilitated the annotation of transcripts using Trinotate, followed by further analysis to obtain the Gene Ontology (GO) annotations. For visualization of the GO terms, the ‘enrichplot’ and ‘ggupset’ packages in R were employed. These steps collectively provided a comprehensive understanding of the functional attributes of the transcriptome data, to understand the underlying biological processes and pathways associated with the studied organisms (**Supplemental Figures S7-S27)**.

### Annotation and genome quality assessments

The *P. bichir* genome annotation was done using the commonly used BRAKER2 (Brůna et al. 2021). To accomplish this, we first used RepeatModeler (Version 5.8.8) (Flynn et al. 2020) to model the repeats in the genome sequence. We then used RepeatMasker (A. F. A. Smit) to mask the repeats found with RepeatModeler and remove them from the genome. The masked genome and the transcriptome aligned using HISAT2 (version 2.1.0) (Kim et al. 2019) were then used to annotate the genome using the Genemark-ET option in BRAKER. The genemark.gtf file was subsequently used by Augustus (Gabriel et al. 2021) to model the proteins.

The completeness of the protein sequences was assessed using BUSCO v. (5.5.0) (Simão et al. 2015; Seppey et al. 2019) with both the Actinopterygii (actinopterygii_odb10, busco.ezlab.org) and vertebrate (vertebrata_odb10, busco.ezlab.org) databases. As polypterids are several hundred million years divergent from all other actinopterygians (Dornburg et al. 2014; Near et al. 2014), the use of the second vertebrate-wide database allowed us to verify similar levels of assembly completeness and mitigate against a potentially teleost-biased ray-finned fish database that may not capture loci in a deeply divergent taxon. We additionally conducted an analysis of synteny between our assembly of the *P. bichir* genome and the previously sequenced *P. senegalus* and *E. calabaricus* genomes using D-Genies (Cabanettes and Klopp 2018).

### Comparative analyses of the Actinopterygian mobilome

To assemble a dataset of TE content across major ray-finned fish lineages, we integrated the results of the repeat analysis of the *P. bichir* genome with other publicly available analyses of transposable elements in ray-finned fishes. For non-teleosts, this captured TE content from two additional polypteriform genomes for *E. calabaricus (Helfenrath et al. 2021)* and *P. senegalus* (Fujito and Nonaka 2012), *Acipenser ruthenus (Du et al. 2020)* as a representative of Chondrostei, as well as *Lepisosteus oculatus* (Braasch et al. 2016), *L. osseus* (Mallik et al. 2023), and *Amia calva* (Thompson et al. 2021) as representative holosteans. These data were integrated with data from 98 teleost genomes previously analyzed for TE content (Reinar et al. 2023), that capture the majority of major teleost lineages including Elopomorpha, Osteoglossomorpha, Otocephala, and a large number of acanthomorph and non-acanthomorph euteleosts. This yielded a dataset of TE content for 105 ray-finned fish genomes. To place this data into a comparative phylogenetic framework, we first obtained a time calibrated phylogeny of all taxa from TimeTree v5 (Kumar et al. 2022). As this tree lacked resolution for acanthomorph lineages, we modified branch lengths to reflect the age estimates from a recent analysis of acanthomorph divergence times based on ultraconserved elements (Ghezelayagh et al. 2022) that is consistent with other published divergence time estimates of this superradiation (Near et al. 2013; Hughes et al. 2018). We additionally modified the topology to reflect the proposed sister relationship between Osteoglossomorpha and Elopomorpha based on genomic and transcriptomic sequence analyses (Hughes et al. 2018; Vialle et al. 2018; Hao et al. 2020; Wcisel et al. 2020).

We used the ggtree and ggtreeExtra packages (Yu 2022) in R 4.3.1 to visualize the distribution of TE abundances (LTR, LINE, SINE, DNA) across the evolutionary history of actinopterygians. We additionally used ggtree in conjunction with phytools (Revell 2012) to visualize genome size and TE content variation between species alongside a likelihood based ancestral state estimation of changes in genome size across the phylogeny conducted in phytools. To assess if changes in genome size were correlated with the changes in the overall abundance of TEs, or changes in the abundances of LTRs, LINEs, SINEs, or DNA elements, we conducted a series of phylogenetic linear regressions using the phylolm package in R. Model fits were assessed by quantification of Akaike Information Criterion (AIC) scores (**Supplemental Table S4**) Regressions were conducted under a Brownian model of trait evolution (**Supplemental Table S6**). To assess whether results were robust to the underlying model of character evolution, analyses were repeated using an Ornstein-Uhlenbeck (OU) model of character evolution (**Supplemental Table S7**). A similar set of phylogenetic regressions was next conducted to assess if increases in the abundance of elements (e.g., LINEs, SINEs, etc) were positively correlated with each other, or if negative correlations exist, allowing us to assess whether elements have antagonistic evolutionary dynamics (**Supplemental Table S8**). Next, we performed a set of regressions assessing whether changes in the abundances of elements were correlated with changes in maximum body size, latitude, or maximum depth of occurrence. Body size and depth data for each species were taken from fishbase using the rfishbase package in R (Boettiger et al. 2012). Latitudinal data was calculated using occurrence data from the Global Biodiversity Information Facility (gbif) using rgbif v3.7.8. We further tested for differences in TE content between teleosts and non-teleosts using a phylogenetic ANOVA in phytools phylANOVA with multiple comparison correction using the Benjamini-Hochberg procedure (Benjamini and Hochberg 1995).

To reconstruct the ancestral mobilome of early ray-finned fishes, we focused on the 25 elements most commonly studied in analyses of ray-finned fish TEs. Presence/absence data was scored for each species and each element and used as input data for a model averaged stochastic character mapping approach (Bollback 2006) implemented in phytools. This approach expands the standard phytools implementation to ancestral state estimation by allowing possible models of character change (e.g., “Equal Rates”, “All Rates Different”) in phytools to contribute to the reconstruction in relation to their Akaike weight. We assessed the degree to which variation in TE content could be explained by evolutionary history through quantification of the phylogenetic signal of overall TE abundance as well as the abundances of each TE type. This was accomplished using the phylosig function in phytools to calculate both Pagel’s λ (Pagel 1999) and Blomberg et al.’s *K* (Blomberg et al. 2003) Values of λ are distributed between zero and 1, with a zero value representing the absence of phylogenetic signal and a value of 1 corresponding to the expectations of Brownian motion on a phylogeny. To assess statistical significance, we compared our empirical λ values to the null expectation that λ = 0 for each trait via a likelihood ratio test. Blomberg’s *K* compliments estimates of λ, with values less than 1 indicating less phylogenetic signal than would be expected given a model of Brownian motion, and value greater than 1 indicating a higher than expected coupling between the distribution of traits and the underlying phylogeny. To assess statistical significance of *K* we compared our empirical *K* values to null distributions of expected *K* values based on 10,000 permutations of each trait on the phylogeny.

To visualize the major axes of variation of the ray-finned fish mobilome, we conducted a phylogenetic principal component analysis (pPCA) using the phyl.pca function in phytools. The resulting PC axes were then used with the phylomorphospace function in phytools to project the phylogeny into the resulting PCA space using ggplot2. Additionally, we conducted a pPCA on the abundances of the subtypes of all TEs based on the amounts derived from the .out files. As preliminary analyses indicated a correlation between elements and overall genome size, pPCAs were repeated on the residuals resulting from a regression of genome size vs TE content, mirroring similar approaches to accounting for traits that covary with another trait (e.g., limb proportions and body size, etc).

## Data Availability

The *P. bichir* genome sequence is available through NCBI Bioproject PRJNA811142. Raw transcriptome reads are available through NCBI under the accession number SRR19537224, SRR19537225, SRR19537226, SRR19537227, SRR19537228, SRR19537229, SRR19537230. Computationally assembled transcriptome sequences are available on NCBI under the accession number GKOV000000000. All the files and code used for TE analysis and visualization are available on Zenodo (DOI: **<u>10.5281/zenodo.10398557</u>)**.

## Acknowledgements

We thank Kent Passingham (NC State University) for assistance with blood collection and the NCBI staff for help with sequence contamination identification. We thank Katerina Zapfe for her invaluable input on figure refinement, and Brandon Turner for helpful insights on transposable elements.

## References

A. F. A. Smit RH&. PG. RepeatMasker. Available from: https://www.repeatmasker.org/webrepeatmaskerhelp.html

Albert JS, Tagliacollo VA, Dagosta F. 2020. Diversification of neotropical freshwater fishes. Annu. Rev. Ecol. Evol. Syst. 51:27–53.

Alonso C, Pérez R, Bazaga P, Herrera CM. 2015. Global DNA cytosine methylation as an evolving trait: phylogenetic signal and correlated evolution with genome size in angiosperms. Front. Genet. 6:4.

Aparicio S, Chapman J, Stupka E, Putnam N, Chia J-M, Dehal P, Christoffels A, Rash S, Hoon S, Smit A, et al. 2002. Whole-genome shotgun assembly and analysis of the genome of Fugu rubripes. Science 297:1301–1310.

Bachmann K, Goin OB, Goin CJ. 1972. The Nuclear DNA of Polypterus palmas. Copeia [Internet] 1972:363. Available from: 10.2307/1442502

Baduel P, Leduque B, Ignace A, Gy I, Gil J Jr, Loudet O, Colot V, Quadrana L. 2021. Genetic and environmental modulation of transposition shapes the evolutionary potential of Arabidopsis thaliana. Genome Biol. 22:138.

Beaulieu JM, Leitch IJ, Patel S, Pendharkar A, Knight CA. 2008. Genome size is a strong predictor of cell size and stomatal density in angiosperms. New Phytol. 179:975–986.

Benjamini Y, Hochberg Y. 1995. Controlling the false discovery rate: A practical and powerful approach to multiple testing. J. R. Stat. Soc. 57:289–300.

Bi X, Wang K, Yang L, Pan H, Jiang H, Wei Q, Fang M, Yu H, Zhu C, Cai Y, et al. 2021. Tracing the genetic footprints of vertebrate landing in non-teleost ray-finned fishes. Cell 184:1377–1391.e14.

Blomberg SP, Garland T Jr, Ives AR. 2003. Testing for phylogenetic signal in comparative data: behavioral traits are more labile. Evolution 57:717–745.

de Boer JG, Yazawa R, Davidson WS, Koop BF. 2007. Bursts and horizontal evolution of DNA transposons in the speciation of pseudotetraploid salmonids. BMC Genomics 8:422.

Boettiger C, Lang DT, Wainwright PC. 2012. rfishbase: exploring, manipulating and visualizing FishBase data from R. J. Fish Biol. 81:2030–2039.

Böhne A, Brunet F, Galiana-Arnoux D, Schultheis C, Volff J-N. 2008. Transposable elements as drivers of genomic and biological diversity in vertebrates. Chromosome Res. 16:203–215.

Bolger AM, Lohse M, Usadel B. 2014. Trimmomatic: a flexible trimmer for Illumina sequence data. Bioinformatics 30:2114–2120.

Bollback JP. 2006. SIMMAP: stochastic character mapping of discrete traits on phylogenies. BMC Bioinformatics 7:88.

Braasch I, Gehrke AR, Smith JJ, Kawasaki K, Manousaki T, Pasquier J, Amores A, Desvignes T, Batzel P, Catchen J, et al. 2016. The spotted gar genome illuminates vertebrate evolution and facilitates human-teleost comparisons. Nat. Genet. 48:427–437.

Britz R, David Johnson G. 2003. On the homology of the posteriormost gill arch in polypterids (Cladistia, Actinopterygii). Zoological Journal of the Linnean Society [Internet] 138:495–503. Available from: 10.1046/j.1096-3642.2003.t01-1-00067.x

Britz R, Johnson GD. 2010. Occipito-vertebral fusion in actinopterygians: conjecture, myth and reality. Part 1: non-teleosts. Origin and Phylogenetic Interrelationships of Teleosts Honoring Gloria Arratia [Internet]. Available from: https://repository.si.edu/bitstream/handle/10088/9748/vz_10Britzand_Johnson.pdf

Brůna T, Hoff KJ, Lomsadze A, Stanke M, Borodovsky M. 2021. BRAKER2: automatic eukaryotic genome annotation with GeneMark-EP+ and AUGUSTUS supported by a protein database. NAR Genom Bioinform 3:lqaa108.

Brunet FG, Roest Crollius H, Paris M, Aury J-M, Gibert P, Jaillon O, Laudet V, Robinson-Rechavi M. 2006. Gene loss and evolutionary rates following whole-genome duplication in teleost fishes. Mol. Biol. Evol. 23:1808–1816.

Cabanettes F, Klopp C. 2018. D-GENIES: dot plot large genomes in an interactive, efficient and simple way. PeerJ 6:e4958.

Capy P, Gasperi G, Biémont C, Bazin C. 2000. Stress and transposable elements: co-evolution or useful parasites? Heredity 85 (Pt 2):101–106.

Carducci F, Barucca M, Canapa A, Carotti E, Biscotti MA. 2020. Mobile Elements in Ray-Finned Fish Genomes. Life [Internet] 10. Available from: 10.3390/life10100221

Chalopin D, Fan S, Simakov O, Meyer A, Schartl M, Volff J-N. 2014. Evolutionary active transposable elements in the genome of the coelacanth. J. Exp. Zool. B Mol. Dev. Evol. 322:322–333.

Chalopin D, Naville M, Plard F, Galiana D, Volff J-N. 2015. Comparative analysis of transposable elements highlights mobilome diversity and evolution in vertebrates. Genome Biol. Evol. 7:567–580.

Cheng P, Huang Y, Du H, Li C, Lv Y, Ruan R, Ye H, Bian C, You X, Xu J, et al. 2019. Draft Genome and Complete Hox-Cluster Characterization of the Sterlet (Acipenser ruthenus). Front. Genet. 10:776.

Crow KD, Wagner GP, SMBE Tri-National Young Investigators. 2006. Proceedings of the SMBE Tri-National Young Investigators’ Workshop 2005. What is the role of genome duplication in the evolution of complexity and diversity? Mol. Biol. Evol. 23:887–892.

Daane JM, Dornburg A, Smits P, MacGuigan DJ, Brent Hawkins M, Near TJ, William Detrich H Iii, Harris MP. 2019. Historical contingency shapes adaptive radiation in Antarctic fishes. Nat Ecol Evol 3:1102–1109.

Davis MP, Holcroft NI, Wiley EO, Sparks JS, Leo Smith W. 2014. Species-specific bioluminescence facilitates speciation in the deep sea. Mar. Biol. 161:1139–1148.

De-Kayne R, Selz OM, Marques DA, Frei D, Seehausen O, Feulner PGD. 2022. Genomic architecture of adaptive radiation and hybridization in Alpine whitefish. Nat. Commun. 13:4479.

Dodsworth S, Chase MW, Kelly LJ, Leitch IJ, Macas J, Novák P, Piednoël M, Weiss-Schneeweiss H, Leitch AR. 2015. Genomic repeat abundances contain phylogenetic signal. Syst. Biol. 64:112–126.

Dornburg A, Federman S, Lamb AD, Jones CD, Near TJ. 2017. Cradles and museums of Antarctic teleost biodiversity. Nat Ecol Evol 1:1379–1384.

Dornburg A, Townsend JP, Friedman M, Near TJ. 2014. Phylogenetic informativeness reconciles ray-finned fish molecular divergence times. BMC Evol. Biol. 14:169.

Du K, Stöck M, Kneitz S, Klopp C, Woltering JM, Adolfi MC, Feron R, Prokopov D, Makunin A, Kichigin I, et al. 2020. The sterlet sturgeon genome sequence and the mechanisms of segmental rediploidization. Nat Ecol Evol 4:841–852.

Fan G, Song Y, Yang L, Huang X, Zhang S, Zhang M, Yang X, Chang Y, Zhang H, Li Y, et al. 2020. Initial data release and announcement of the 10,000 Fish Genomes Project (Fish10K). Gigascience [Internet] 9. Available from: 10.1093/gigascience/giaa080

Flynn JM, Hubley R, Goubert C, Rosen J, Clark AG, Feschotte C, Smit AF. 2020. RepeatModeler2 for automated genomic discovery of transposable element families. Proc. Natl. Acad. Sci. U. S. A. 117:9451–9457.

Fujino K, Hashida S-N, Ogawa T, Natsume T, Uchiyama T, Mikami T, Kishima Y. 2011. Temperature controls nuclear import of Tam3 transposase in Antirrhinum. Plant J. 65:146–155.

Fujito NT, Nonaka M. 2012. Highly divergent dimorphic alleles of the proteasome subunit beta type-8 (PSMB8) gene of the bichir Polypterus senegalus: implication for evolution of the PSMB8 gene of jawed vertebrates. Immunogenetics 64:447–453.

Gabriel L, Hoff KJ, Brůna T, Borodovsky M, Stanke M. 2021. TSEBRA: transcript selector for BRAKER. BMC Bioinformatics 22:566.

Galbraith JD, Ludington AJ, Sanders KL, Amos TG, Thomson VA, Enosi Tuipulotu D, Dunstan N, Edwards RJ, Suh A, Adelson DL. 2022. Horizontal Transposon Transfer and Its Implications for the Ancestral Ecology of Hydrophiine Snakes. Genes [Internet] 13. Available from: 10.3390/genes13020217

Ghezelayagh A, Harrington RC, Burress ED, Campbell MA, Buckner JC, Chakrabarty P, Glass JR, McCraney WT, Unmack PJ, Thacker CE, et al. 2022. Prolonged morphological expansion of spiny-rayed fishes following the end-Cretaceous. Nat Ecol Evol 6:1211–1220.

Glasauer SMK, Neuhauss SCF. 2014. Whole-genome duplication in teleost fishes and its evolutionary consequences. Mol. Genet. Genomics 289:1045–1060.

Grabherr MG, Haas BJ, Yassour M, Levin JZ, Thompson DA, Amit I, Adiconis X, Fan L, Raychowdhury R, Zeng Q, et al. 2011. Full-length transcriptome assembly from RNA-Seq data without a reference genome. Nat. Biotechnol. 29:644–652.

Hao S, Han K, Meng L, Huang X, Shi C, Zhang M, Wang Y, Liu Q, Zhang Y, Seim I, et al. 2020. Three genomes of Osteoglossidae shed light on ancient teleost evolution. bioRxiv [Internet]. Available from: http://biorxiv.org/lookup/doi/10.1101/2020.01.19.911958

Helfenrath K, Sauer M, Kamga M, Wisniewsky M, Burmester T, Fabrizius A. 2021. The More, the Merrier? Multiple Myoglobin Genes in Fish Species, Especially in Gray Bichir (Polypterus senegalus) and Reedfish (Erpetoichthys calabaricus). Genome Biol. Evol. [Internet] 13. Available from: 10.1093/gbe/evab078

Hickman AB, Dyda F. 2015. Mechanisms of DNA Transposition. Microbiol Spectr 3:MDNA3–MDNA0034–2014.

Hua-Van A, Le Rouzic A, Boutin TS, Filée J, Capy P. 2011. The struggle for life of the genome’s selfish architects. Biol. Direct 6:19.

Hughes LC, Ortí G, Huang Y, Sun Y, Baldwin CC, Thompson AW, Arcila D, Betancur-R R, Li C, Becker L, et al. 2018. Comprehensive phylogeny of ray-finned fishes (Actinopterygii) based on transcriptomic and genomic data. Proc. Natl. Acad. Sci. U. S. A. 115:6249–6254.

Ivancevic AM, Kortschak RD, Bertozzi T, Adelson DL. 2016. LINEs between Species: Evolutionary Dynamics of LINE-1 Retrotransposons across the Eukaryotic Tree of Life. Genome Biol. Evol. 8:3301–3322.

Jiang S, Cai D, Sun Y, Teng Y. 2016. Isolation and characterization of putative functional long terminal repeat retrotransposons in the Pyrus genome. Mob. DNA 7:1.

Kapitonov VV, Jurka J. 2007. Chapaev-a novel superfamily of DNA transposons. Repbase Reports 7:774–781.

Kelleher ES, Barbash DA, Blumenstiel JP. 2020. Taming the Turmoil Within: New Insights on the Containment of Transposable Elements. Trends Genet. 36:474–489.

Kim D, Paggi JM, Park C, Bennett C, Salzberg SL. 2019. Graph-based genome alignment and genotyping with HISAT2 and HISAT-genotype. Nat. Biotechnol. 37:907–915.

Kimura Y, Nikaido M. 2021. Conserved Keratin Gene Clusters in Ancient Fish: an Evolutionary Seed for Terrestrial Adaptation. Genomics 113:1120–1128.

Kordis D, Lovsin N, Gubensek F. 2006. Phylogenomic analysis of the L1 retrotransposons in Deuterostomia. Syst. Biol. 55:886–901.

Krysanov EY, Golubtsov AS. 2014. Karyotypes of four fish species from the Nile and Omo-Turkana basins in Ethiopia. J. Ichthyol. 54:889–892.

Kumar S, Suleski M, Craig JM, Kasprowicz AE, Sanderford M, Li M, Stecher G, Hedges SB. 2022. TimeTree 5: An Expanded Resource for Species Divergence Times. Mol. Biol. Evol. [Internet] 39. Available from: 10.1093/molbev/msac174

Lehmann R, Kovařík A, Ocalewicz K, Kirtiklis L, Zuccolo A, Tegner JN, Wanzenböck J, Bernatchez L, Lamatsch DK, Symonová R. 2021. DNA Transposon Expansion is Associated with Genome Size Increase in Mudminnows. Genome Biol. Evol. [Internet] 13. Available from: 10.1093/gbe/evab228

Lieberman-Aiden E, van Berkum NL, Williams L, Imakaev M, Ragoczy T, Telling A, Amit I, Lajoie BR, Sabo PJ, Dorschner MO, et al. 2009. Comprehensive mapping of long-range interactions reveals folding principles of the human genome. Science 326:289–293.

Lo R, Dougan KE, Chen Y, Shah S, Bhattacharya D, Chan CX. 2022. Alignment-Free Analysis of Whole-Genome Sequences From Symbiodiniaceae Reveals Different Phylogenetic Signals in Distinct Regions. Front. Plant Sci. 13:815714.

Lynch M. 2007. The frailty of adaptive hypotheses for the origins of organismal complexity. Proc. Natl. Acad. Sci. U. S. A. 104 Suppl 1:8597–8604.

Mallik R, Carlson KB, Wcisel DJ, Fisk M, Yoder JA, Dornburg A. 2023. A chromosome-level genome assembly of longnose gar, Lepisosteus osseus. G3 [Internet] 13. Available from: 10.1093/g3journal/jkad095

Manni M, Berkeley MR, Seppey M, Simão FA, Zdobnov EM. 2021. BUSCO Update: Novel and Streamlined Workflows along with Broader and Deeper Phylogenetic Coverage for Scoring of Eukaryotic, Prokaryotic, and Viral Genomes. Mol. Biol. Evol. 38:4647–4654.

Martinez-Gutierrez CA, Aylward FO. 2022. Genome size distributions in bacteria and archaea are strongly linked to evolutionary history at broad phylogenetic scales. PLoS Genet. 18:e1010220.

Matzke MA, Matzke AJ. 1998. Polyploidy and transposons. Trends Ecol. Evol. 13:241.

McClintock B. 1984. The significance of responses of the genome to challenge. Science 226:792–801.

Mikami M, Ineno T, Thompson AW, Braasch I, Ishiyama M, Kawasaki K. 2022. Convergent losses of SCPP genes and ganoid scales among non-teleost actinopterygians. Gene 811:146091.

Miller EC, Martinez CM, Friedman ST, Wainwright PC, Price SA, Tornabene L. 2022. Alternating regimes of shallow and deep-sea diversification explain a species-richness paradox in marine fishes. Proc. Natl. Acad. Sci. U. S. A. 119:e2123544119.

Morescalchi MA, Liguori I, Rocco L, Stingo V. 2007. Karyotypic characterization and genomic organization of the 5S rDNA in Erpetoichthys calabaricus (Osteichthyes, Polypteridae). Genetica 131:209–216.

Moritz T, Britz R. Revision of the extant Polypteridae (Actinopterygii: Cladistia). Ichthyol. Explor. Freshw. [Internet]. Available from: 10.23788/IEF-1094

Moriyama Y, Koshiba-Takeuchi K. 2018. Significance of whole-genome duplications on the emergence of evolutionary novelties. Brief. Funct. Genomics 17:329–338.

Muñoz-López M, García-Pérez JL. 2010. DNA transposons: nature and applications in genomics. Curr. Genomics 11:115–128.

Naville M, Henriet S, Warren I, Sumic S, Reeve M, Volff J-N, Chourrout D. 2019. Massive Changes of Genome Size Driven by Expansions of Non-autonomous Transposable Elements. Curr. Biol. 29:1161–1168.e6.

Near TJ, Dornburg A, Eytan RI, Keck BP, Smith WL, Kuhn KL, Moore JA, Price SA, Burbrink FT, Friedman M, et al. 2013. Phylogeny and tempo of diversification in the superradiation of spiny-rayed fishes. Proc. Natl. Acad. Sci. U. S. A. 110:12738–12743.

Near TJ, Dornburg A, Kuhn KL, Eastman JT, Pennington JN, Patarnello T, Zane L, Fernández DA, Jones CD. 2012. Ancient climate change, antifreeze, and the evolutionary diversification of Antarctic fishes. Proc. Natl. Acad. Sci. U. S. A. 109:3434–3439.

Near TJ, Dornburg A, Tokita M, Suzuki D, Brandley MC, Friedman M. 2014. Boom and bust: ancient and recent diversification in bichirs (Polypteridae: Actinopterygii), a relictual lineage of ray-finned fishes. Evolution 68:1014–1026.

Niu X-M, Xu Y-C, Li Z-W, Bian Y-T, Hou X-H, Chen J-F, Zou Y-P, Jiang J, Wu Q, Ge S, et al. 2019. Transposable elements drive rapid phenotypic variation in. Proc. Natl. Acad. Sci. U. S. A. 116:6908–6913.

Novick PA, Smith JD, Floumanhaft M, Ray DA, Boissinot S. 2011. The evolution and diversity of DNA transposons in the genome of the Lizard Anolis carolinensis. Genome Biol. Evol. 3:1–14.

Olmo E, Capriglione T, Odierna G. 1989. Genome size evolution in vertebrates: trends and constraints. Comp. Biochem. Physiol. B 92:447–453.

Pagel M. 1999. Inferring the historical patterns of biological evolution. Nature 401:877–884.

Papon N, Lasserre-Zuber P, Rimbert H, De Oliveira R, Paux E, Choulet F. 2023. All families of transposable elements were active in the recent wheat genome evolution and polyploidy had no impact on their activity. Plant Genome:e20347.

Parisod C, Alix K, Just J, Petit M, Sarilar V, Mhiri C, Ainouche M, Chalhoub B, Grandbastien M-A. 2010. Impact of transposable elements on the organization and function of allopolyploid genomes. New Phytol. 186:37–45.

Parker E, Zapfe KL, Yadav J, Frédérich B, Jones CD, Economo EP, Federman S, Near TJ, Dornburg A. 2022. Periodic Environmental Disturbance Drives Repeated Ecomorphological Diversification in an Adaptive Radiation of Antarctic Fishes. Am. Nat. 200:E221–E236.

Paulat NS, Storer JM, Moreno-Santillán DD, Osmanski AB, Sullivan KAM, Grimshaw JR, Korstian J, Halsey M, Garcia CJ, Crookshanks C, et al. 2023. Chiropterans Are a Hotspot for Horizontal Transfer of DNA Transposons in Mammalia. Mol. Biol. Evol. [Internet] 40. Available from: 10.1093/molbev/msad092

Piskurek O, Jackson DJ. 2011. Tracking the ancestry of a deeply conserved eumetazoan SINE domain. Mol. Biol. Evol. 28:2727–2730.

Putnam NH, O’Connell BL, Stites JC, Rice BJ, Blanchette M, Calef R, Troll CJ, Fields A, Hartley PD, Sugnet CW, et al. 2016. Chromosome-scale shotgun assembly using an in vitro method for long-range linkage. Genome Research [Internet] 26:342–350. Available from: 10.1101/gr.193474.115

Randhawa SS, Pawar R. 2021. Fish genomes: Sequencing trends, taxonomy and influence of taxonomy on genome attributes. J. Appl. Ichthyol. 37:553–562.

Rech GE, Radío S, Guirao-Rico S, Aguilera L, Horvath V, Green L, Lindstadt H, Jamilloux V, Quesneville H, González J. 2022. Population-scale long-read sequencing uncovers transposable elements associated with gene expression variation and adaptive signatures in Drosophila. Nat. Commun. 13:1948.

Reinar WB, Tørresen OK, Nederbragt AJ, Matschiner M, Jentoft S, Jakobsen KS. 2023. Teleost genomic repeat landscapes in light of diversification rates and ecology. Mob. DNA 14:14.

Reiss D, Mialdea G, Miele V, de Vienne DM, Peccoud J, Gilbert C, Duret L, Charlat S. 2019. Global survey of mobile DNA horizontal transfer in arthropods reveals Lepidoptera as a prime hotspot. PLoS Genet. 15:e1007965.

Revell LJ. 2012. phytools: an R package for phylogenetic comparative biology (and other things). Methods Ecol. Evol. 3:217–223.

Revell LJ. 2023. phytools 2.0: An updated R ecosystem for phylogenetic comparative methods (and other things). bioRxiv [Internet]:2023.03.08.531791. Available from: https://www.biorxiv.org/content/10.1101/2023.03.08.531791v2.abstract

Ricci M, Peona V, Guichard E, Taccioli C, Boattini A. 2018. Transposable Elements Activity is Positively Related to Rate of Speciation in Mammals. J. Mol. Evol. 86:303–310.

Rodriguez F, Arkhipova IR. 2018. Transposable elements and polyploid evolution in animals. Curr. Opin. Genet. Dev. 49:115–123.

Senft AD, Macfarlan TS. 2021. Transposable elements shape the evolution of mammalian development. Nat. Rev. Genet. 22:691–711.

Seppey M, Manni M, Zdobnov EM. 2019. BUSCO: Assessing Genome Assembly and Annotation Completeness. Methods Mol. Biol. 1962:227–245.

Serrato-Capuchina A, Matute DR. 2018. The Role of Transposable Elements in Speciation. Genes [Internet] 9. Available from: 10.3390/genes9050254

Sessegolo C, Burlet N, Haudry A. 2016. Strong phylogenetic inertia on genome size and transposable element content among 26 species of flies. Biol. Lett. [Internet] 12. Available from: 10.1098/rsbl.2016.0407

Simão FA, Waterhouse RM, Ioannidis P, Kriventseva EV, Zdobnov EM. 2015. BUSCO: assessing genome assembly and annotation completeness with single-copy orthologs. Bioinformatics 31:3210–3212.

Sotero-Caio CG, Platt RN 2nd, Suh A, Ray DA. 2017. Evolution and Diversity of Transposable Elements in Vertebrate Genomes. Genome Biol. Evol. 9:161–177.

Suzuki D, Brandley MC, Tokita M. 2010. The mitochondrial phylogeny of an ancient lineage of ray-finned fishes (Polypteridae) with implications for the evolution of body elongation, pelvic fin loss, and craniofacial morphology in Osteichthyes. BMC Evol. Biol. 10:209.

Tambones IL, Haudry A, Simão MC, Carareto CMA. 2019. High frequency of horizontal transfer in Jockey families (LINE order) of drosophilids. Mob. DNA 10:43.

Thompson AW, Hawkins MB, Parey E, Wcisel DJ, Ota T, Kawasaki K, Funk E, Losilla M, Fitch OE, Pan Q, et al. 2021. The bowfin genome illuminates the developmental evolution of ray-finned fishes. Nat. Genet. 53:1373–1384.

Torres DE, Thomma BPHJ, Seidl MF. 2021. Transposable Elements Contribute to Genome Dynamics and Gene Expression Variation in the Fungal Plant Pathogen Verticillium dahliae. Genome Biol. Evol. [Internet] 13. Available from: 10.1093/gbe/evab135

Turner BA, Miorin TR, Stewart NB, Reid RW, Moore CC, Rogers RL. 2021. Chromosomal rearrangements as a source of local adaptation in island Drosophila. Available from: https://arxiv.org/abs/2109.09801

Vervoort A. 1980. Karyotypes and nuclear DNA contents of Polypteridae (Osteichthyes). Experientia 36:646–647.

Vialle RA, de Souza JES, Lopes K de P, Teixeira DG, Alves Sobrinho P de A, Ribeiro-Dos-Santos AM, Furtado C, Sakamoto T, Oliveira Silva FA, Herculano Corrêa de Oliveira E, et al. 2018. Whole Genome Sequencing of the Pirarucu (Arapaima gigas) Supports Independent Emergence of Major Teleost Clades. Genome Biol. Evol. 10:2366–2379.

Volff J-N, Bouneau L, Ozouf-Costaz C, Fischer C. 2003. Diversity of retrotransposable elements in compact pufferfish genomes. Trends Genet. 19:674–678.

Wcisel DJ, Howard JT 3rd, Yoder JA, Dornburg A. 2020. Transcriptome Ortholog Alignment Sequence Tools (TOAST) for phylogenomic dataset assembly. BMC Evol. Biol. 20:41.

Wen H, Luo T, Wang Y, Wang S, Liu T, Xiao N, Zhou J. 2022. Molecular phylogeny and historical biogeography of the cave fish genus Sinocyclocheilus (Cypriniformes: Cyprinidae) in southwest China. Integr. Zool. 17:311–325.

Wickham H. 2016. ggplot2: Elegant Graphics for Data Analysis. Available from: https://ggplot2.tidyverse.org

Wong WY, Simakov O, Bridge DM, Cartwright P, Bellantuono AJ, Kuhn A, Holstein TW, David CN, Steele RE, Martínez DE. 2019. Expansion of a single transposable element family is associated with genome-size increase and radiation in the genus. Proc. Natl. Acad. Sci. U. S. A. 116:22915–22917.

Wright JJ, Bruce SA, Sinopoli DA, Palumbo JR, Stewart DJ. 2022. Phylogenomic analysis of the bowfin (Amia calva) reveals unrecognized species diversity in a living fossil lineage. Sci. Rep. 12:16514.

Yu G. 2022. Data Integration, Manipulation and Visualization of Phylogenetic Trees. CRC Press

Zhang H-H, Feschotte C, Han M-J, Zhang Z. 2014. Recurrent horizontal transfers of Chapaev transposons in diverse invertebrate and vertebrate animals. Genome Biol. Evol. 6:1375–1386.

Zhang H-H, Peccoud J, Xu M-R-X, Zhang X-G, Gilbert C. 2020. Horizontal transfer and evolution of transposable elements in vertebrates. Nat. Commun. 11:1362.

Zhang Z, Sakuma A, Kuraku S, Nikaido M. 2021. Evolutionary Transition From Water to Land in Vertebrates Illuminated by Basal Ray-Finned Fish Vomeronasal Type 2 Receptor (OlfC) Genes. preprint [Internet]. Available from: 10.21203/rs.3.rs-504504/v1

