## Supplemental Information for "Investigating the impact of whole genome duplication on transposable element evolution in ray-finned fishes"

| | |
|---|---|
| J.A. Yoder: | 0000-0002-6083-1311 |
| D.J. Wcisel: | 0000-0001-6858-643X |
| T.J. Near | 0000-0002-7398-6670 |
| R. Mallik: | 0000-0002-6594-1835 |
| A. Dornburg: | 0000-0003-0863-2283 |

*** Corresponding author at:**

**Contents:**


| | | |
|---|---|---|
| Supplemental Figure S1 | Busco analysis comparison of *Polypterus bichir* and *Polypterus senegalus*. | 1 |
| Supplemental Figure S2 | Synteny plots | 2 |
| Supplemental Figure S3 | Biplots of the phylomorphospaces corresponding to Figure 5 | 2 |
| Supplemental Figure S4 | Labelled plots of results of a phylogenetic PCA for Figure 5A | 3 |
| Supplemental Figure S5 | Labelled plots of results of a phylogenetic PCA for Figure 5B | 3 |
| Supplemental Figure S6 | Labelled plots of results of a phylogenetic PCA for Figure 5C | 4 |
| Supplemental Figure S7 | Summary of Gene Ontology *Biological Processes* analysis from the *Polypterus bichir* gill transcriptome. | 5 |
| Supplemental Figure S8 | Summary of Gene Ontology *Molecular Function* analysis from the *Polypterus bichir* gill transcriptome. | 6 |
| Supplemental Figure S9 | Summary of Gene Ontology Cellular Components analysis from the *Polypterus bichir* gill transcriptome | 7 |
| Supplemental Figure S10 | Summary of Gene Ontology *Cellular Components* analysis from the *Polypterus bichir* kidney transcriptome. | 8 |
| Supplemental Figure S11 | Summary of Gene Ontology Biological Processes analysis from the *Polypterus bichir* kidney transcriptome | 9 |
| Supplemental Figure S12 | Summary of Gene Ontology *Molecular Functions* analysis from the *Polypterus bichir* kidney transcriptome. | 10 |
| Supplemental Figure S13 | Summary of Gene Ontology *Biological Processes* analysis from the *Polypterus bichir* liver transcriptome | 11 |
| Supplemental Figure S14 | Summary of Gene Ontology Molecular Functions analysis from the Polypterus bichir liver transcriptome | 12 |
| Supplemental Figure S15 | Summary of Gene Ontology *Cellular Components* analysis from the *Polypterus bichir* liver transcriptome | 13 |
| Supplemental Figure S16 | Summary of Gene Ontology *Biological Processes* analysis from the *Polypterus bichir* spleen transcriptome | 14 |
| Supplemental Figure S17 | Summary of Gene Ontology *Molecular Function* analysis from the *Polypterus bichir* spleen transcriptome | 15 |
| Supplemental Figure S18 | Summary of Gene Ontology *Cellular Components* analysis from the *Polypterus bichir* spleen transcriptome | 16 |

| | | |
|---|---|---|
| Supplemental Figure S19 | Summary of Gene Ontology *Biological Processes* analysis from the *Polypterus bichir* gut transcriptome | 17 |
| Supplemental Figure S20 | Summary of Gene Ontology *Molecular Functions* analysis from the *Polypterus bichir* gut transcriptome. | 18 |
| Supplemental Figure S22 | Summary of Gene Ontology *Molecular Functions* analysis from the *Polypterus bichir* heart transcriptome | 19 |
| Supplemental Figure S23 | Summary of Gene Ontology *Cellular Components* analysis from the *Polypterus bichir* heart transcriptome. | 21 |
| Supplemental Figure S24 | Summary of Gene Ontology *Biological Processes* analysis from the *Polypterus bichir* heart transcriptome. | 22 |
| Supplemental Figure S25 | Summary of Gene Ontology *Biological Processes* analysis from the *Polypterus bichir* eye transcriptome. | 23 |
| Supplemental Figure S26 | Summary of Gene Ontology *Cellular Components* analysis from the *Polypterus bichir* eye transcriptome. | 24 |
| Supplemental Figure S27 | Summary of Gene Ontology *Cellular Components* analysis from the *Polypterus bichir* eye transcriptome | 25 |
| Supplemental Table S1 | Comparative genome assembly metrics of *Polypterus bichir* and related species | 26 |
| Supplemental Table S2 | Comparative analysis of genome completeness of *Polypterus bichir* and related species | 26 |
| Supplemental Table S3 | Phylogenetic Signal and Variance Analysis Using Pagel's Lambda, Blomberg's K, and ANOVA | 27 |
| Supplemental Table S4 | Evaluation of model fit for different TE types, genome size, and habitat characteristics under Brownian motion and Ornstein-Uhlenbeck (OU) models | 27 |
| Supplemental Table S5 | AIC scores from phylogenetic linear models evaluating each type of transposable element (TE) | 28 |
| Supplemental Table S6 | Phylogenetic linear models depict the correlation between each type of transposable element (TE) percentages and genome size, under the Brownian motion correlation model. | 28 |
| Supplemental Table S7 | Results of phylogenetic linear models evaluating the correlation between each type of transposable element (TE) percentages and genome size, under the OU correlation model. | 29 |
| Supplemental Table S8 | Phylogenetic linear models' outcomes depict the correlation between percentages of each transposable element (TE) type | 29 |

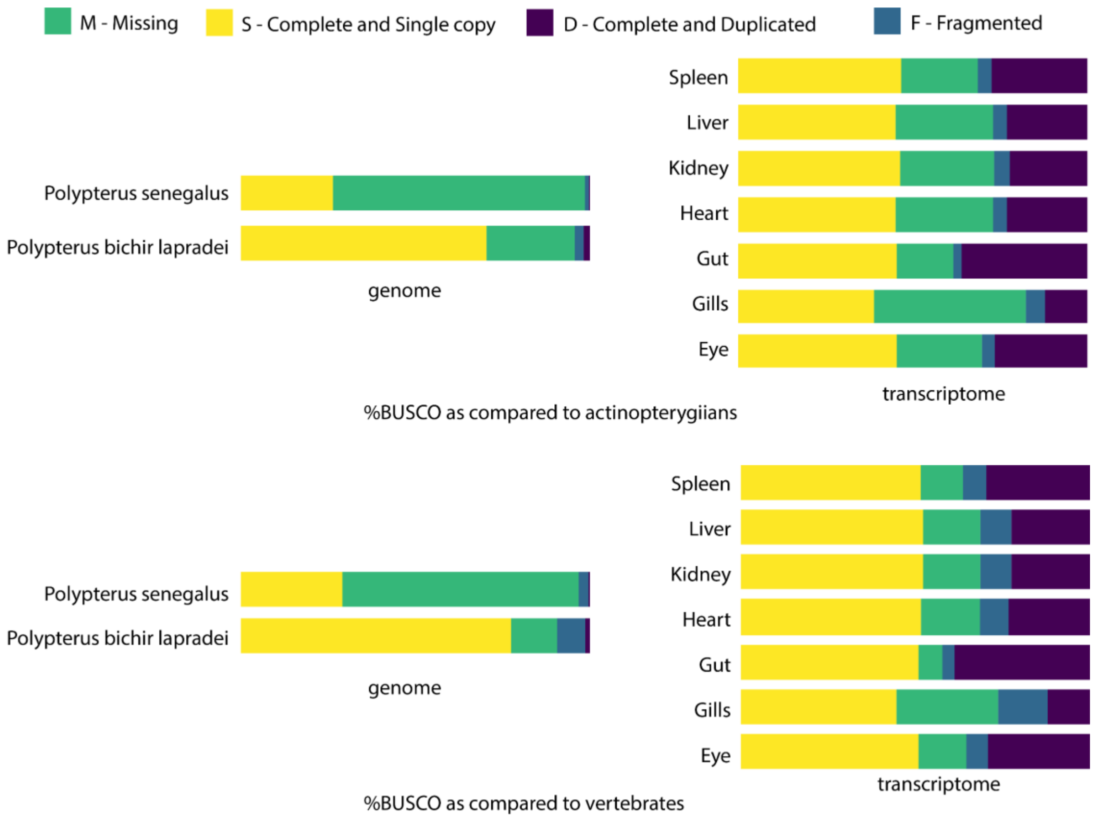


**Supplemental Figure S1.** BUSCO analysis comparing the *Polypterus bichir* and *Polypterus senegalus* reference genomes and our transcripts for *Polypterus bichir* from various tissues. BUSCO scores for the *Polypterus bichir* reference genome were higher than those of the *Polypterus senegalus* reference, indicating the higher completeness and quality of the *Polypterus bichir* genome assembly. The upper panel depicts the loci captured when using the actinopterygian BUSCO reference set, the lower panel indicates the loci captured when utilizing the vertebrate reference set.

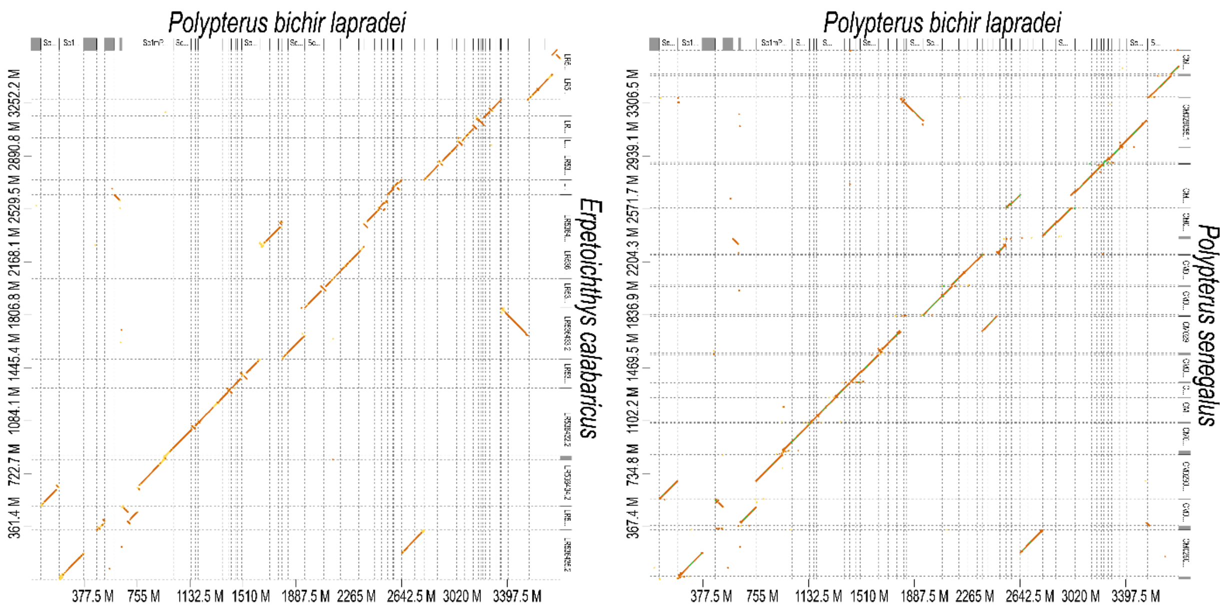


**Supplemental Figure S2:** D-genies plot showing synteny between the *Polypterus bichir* and *Polypterus senegalus* reference genomes (left) and between the *Polypterus bichir* and *Erpetoichthys calabaricus* reference genomes (right).

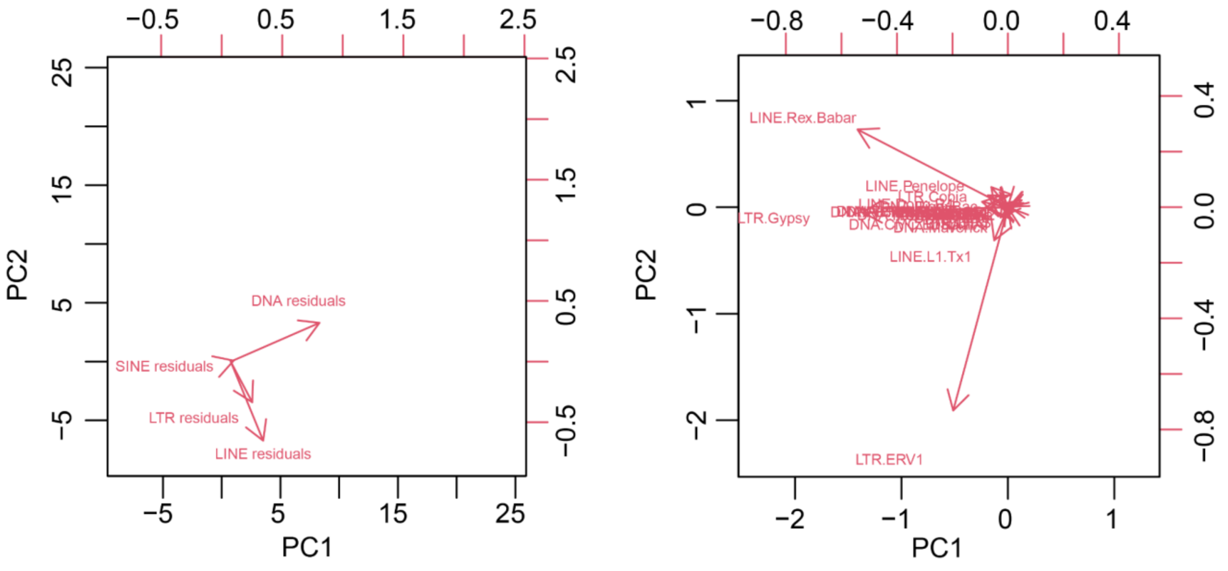


**Supplemental Figure S3:** Biplots depicting the loading of the variables in the phylogenetic PCA spaces depicted in Figure 5

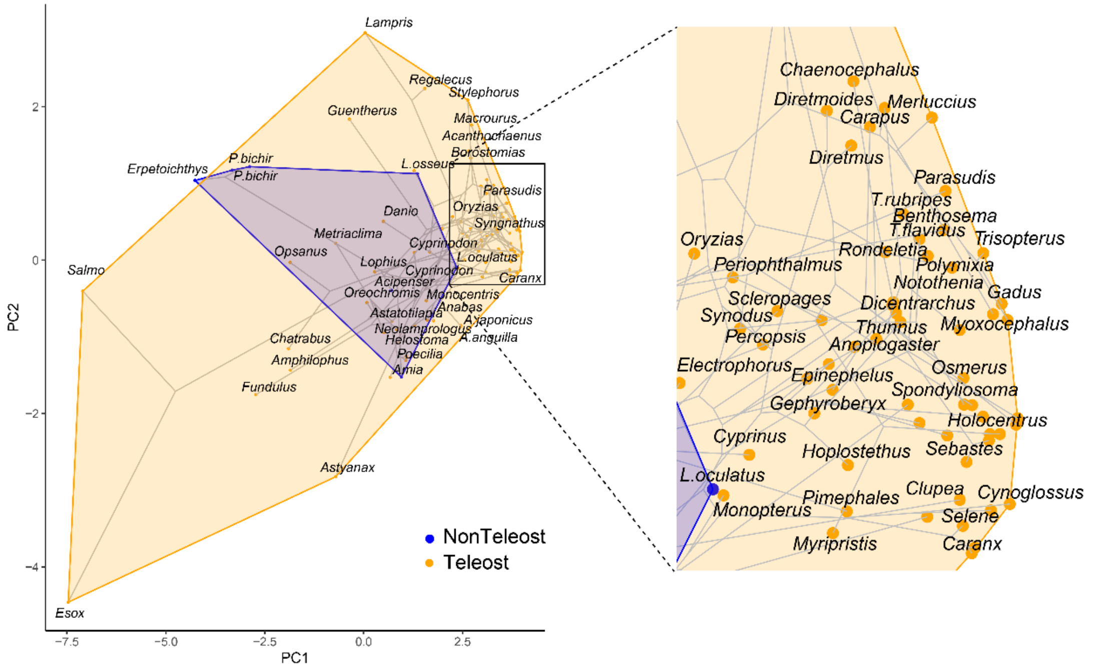


**Supplemental Figure S4:** Labelled plots of results of a phylogenetic PCA on the abundances of LTRs, DNA transposons, LINEs, and SINEs that project the phylogeny and points onto a 2D plot of PC1 and PC2. Blue indicates non-teleost actinopterygians, orange indicates teleosts.

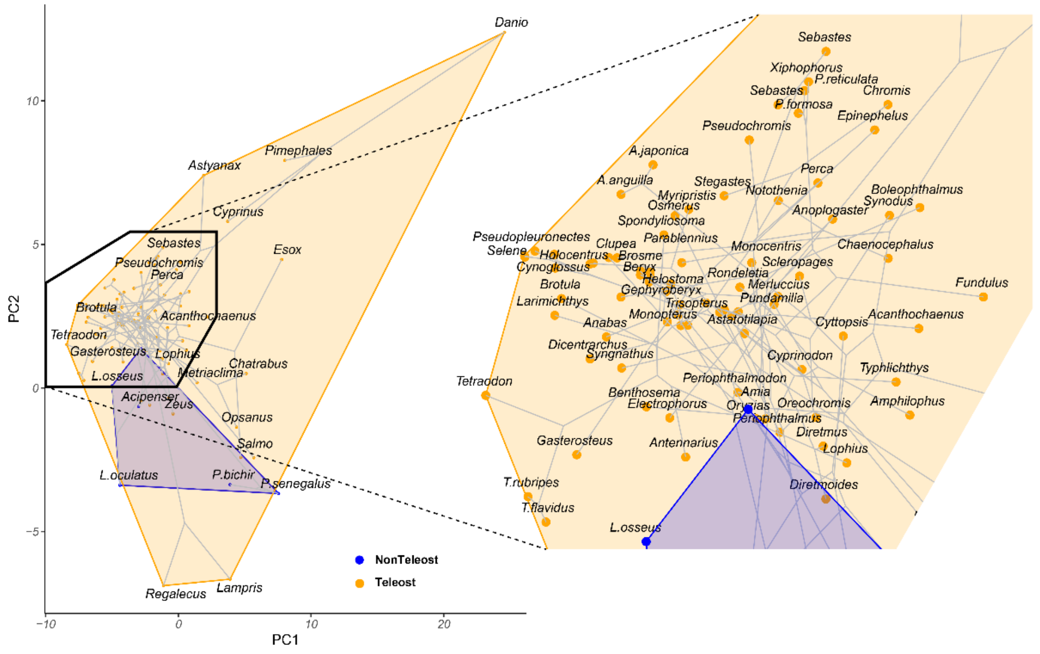


**Supplemental Figure S5:** Labelled plots of phylomorphospace depicting the resulting space based on the residual variation following linear regression between major TE classes and genome size. Blue indicates non-teleost actinopterygians, orange indicates teleosts.

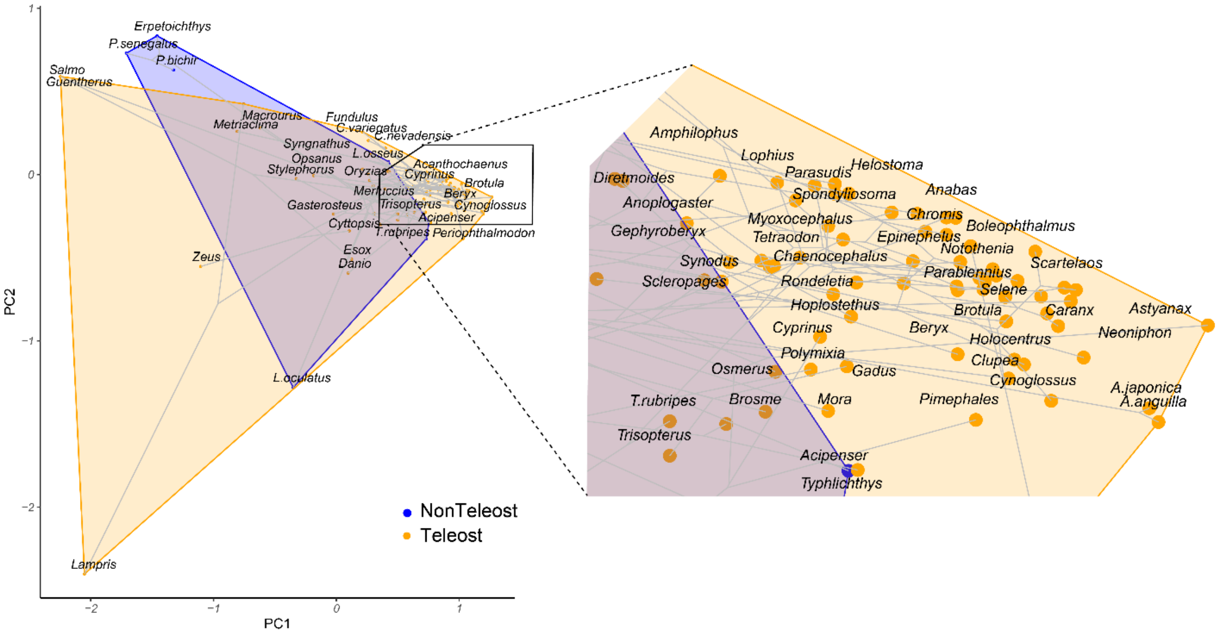


**Supplemental Figure S6:** Labelled plots depicting the results of a PCA on the residual variation of the abundances of all 25 superfamilies accounting for differences in genome size, consisting of 13 DNA transposons, 7 LINEs, and 5 LTR subfamilies. Blue indicates non-teleost actinopterygians, orange indicates teleosts.

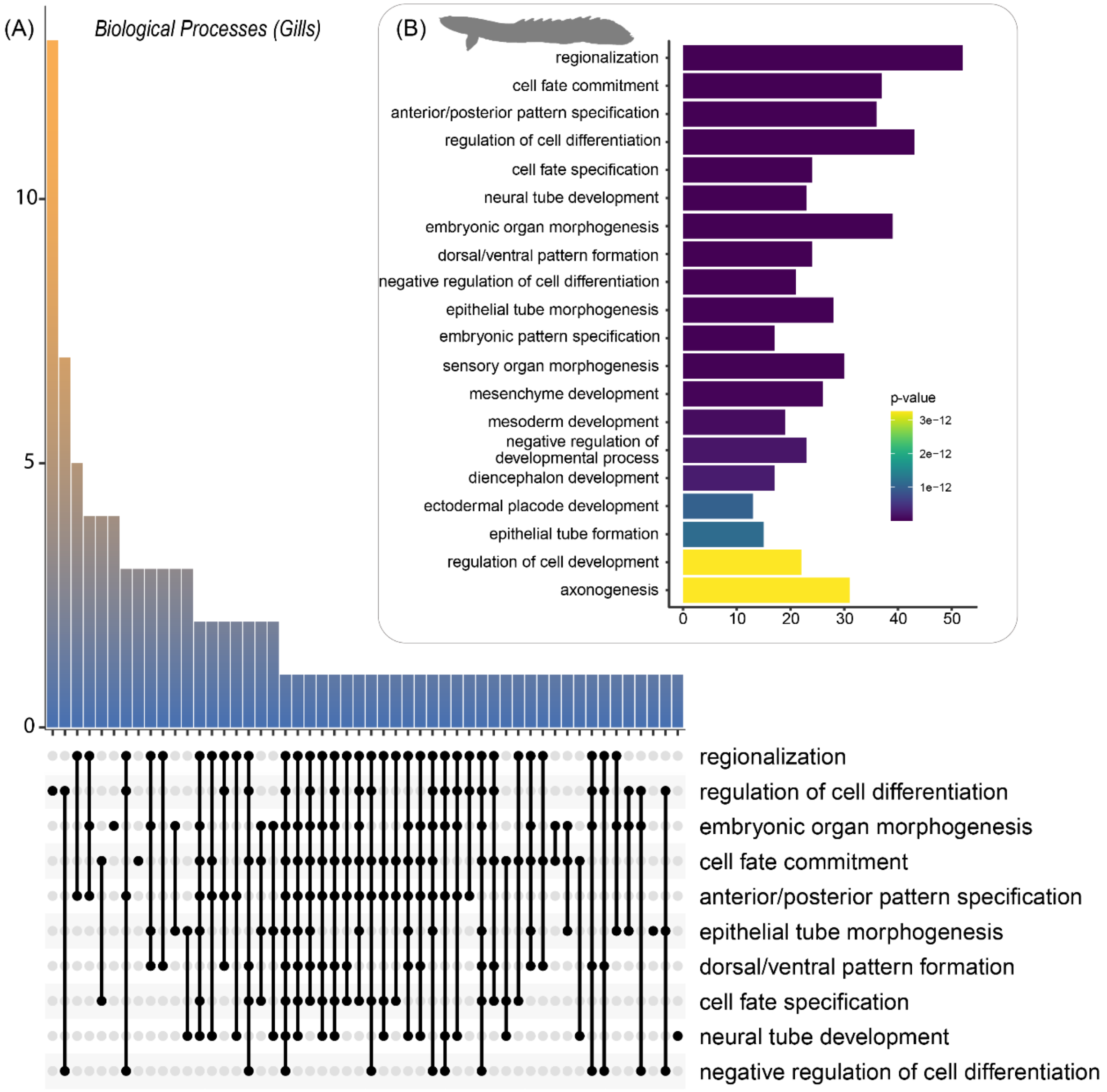


**Supplemental Figure S7:** Summary of Gene Ontology *Biological Processes* analysis from the *Polypterus bichir* gill transcriptome.
Predicted proteins from the transcriptome were used as inputs to assess (A) processes and their intersections and (B) most common terms.

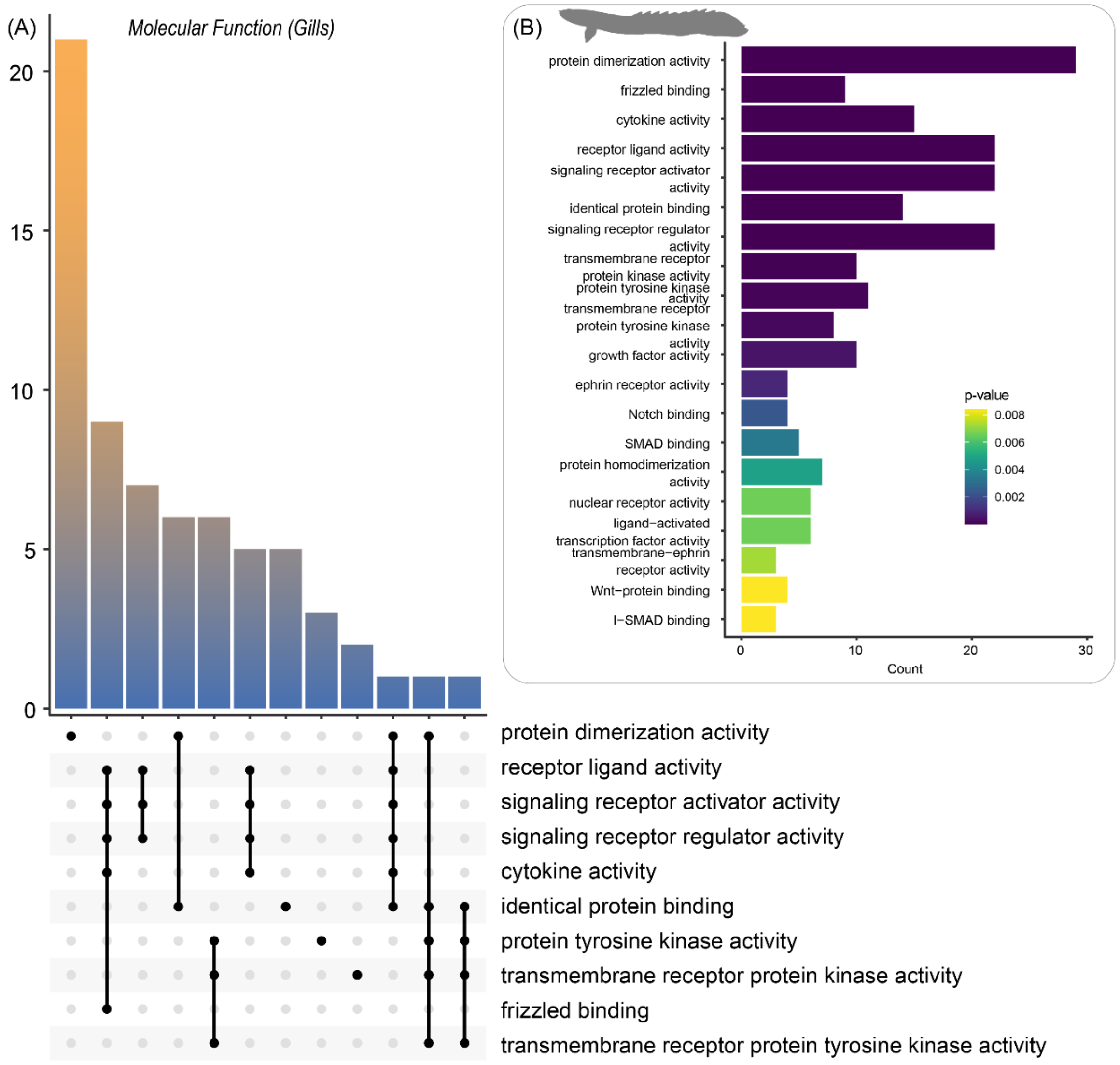


**Supplemental Figure S8:** Summary of Gene Ontology *Molecular Function* analysis from the *Polypterus bichir* gill transcriptome.
Predicted proteins from the transcriptome were used as inputs to assess (A) processes and their intersections and (B) most common terms.

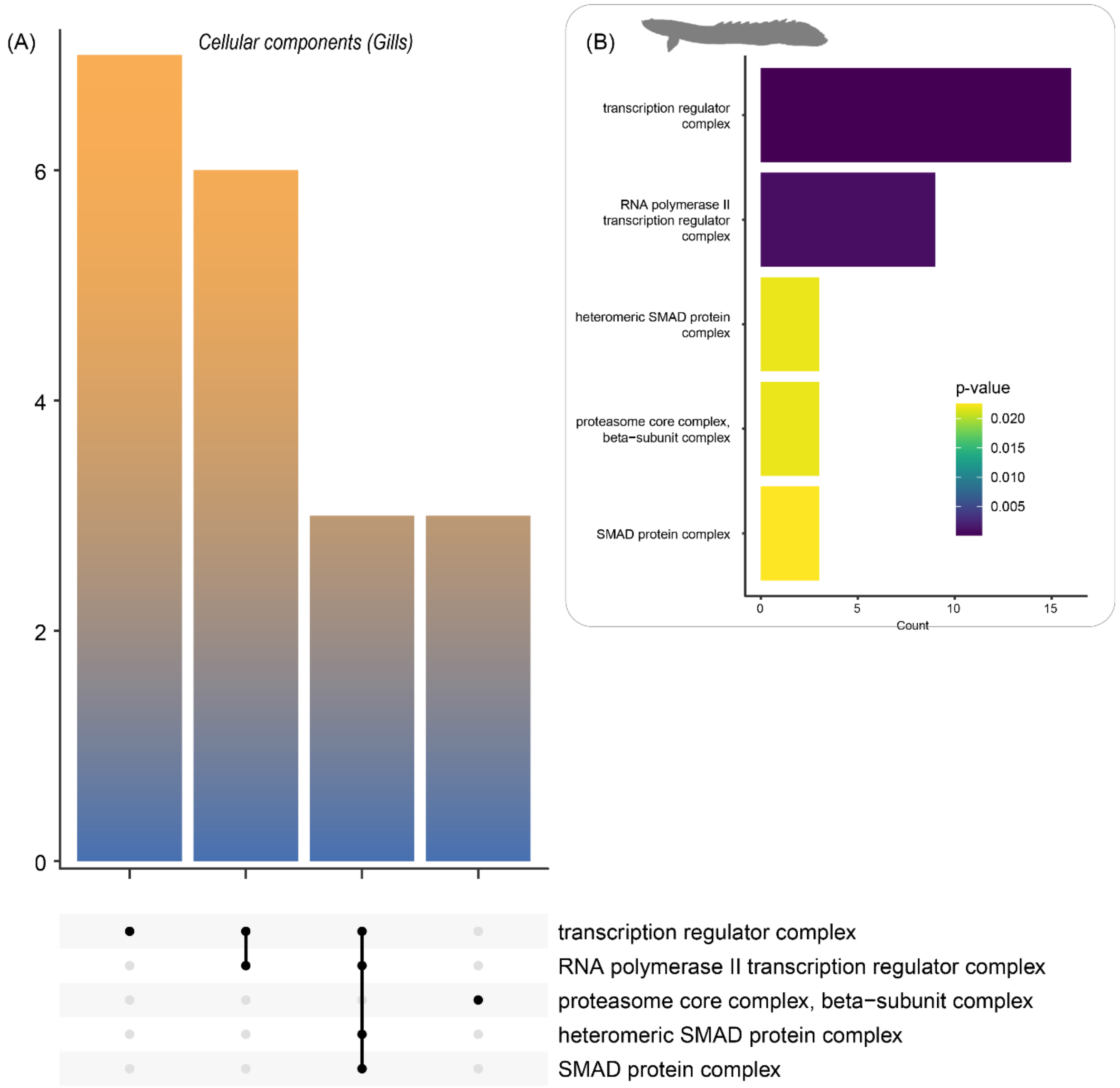


**Supplemental Figure S9:** Summary of Gene Ontology *Cellular Components* analysis from the *Polypterus bichir* gill transcriptome.
Predicted proteins from the transcriptome were used as inputs to assess (A) processes and their intersections and (B) most common terms.

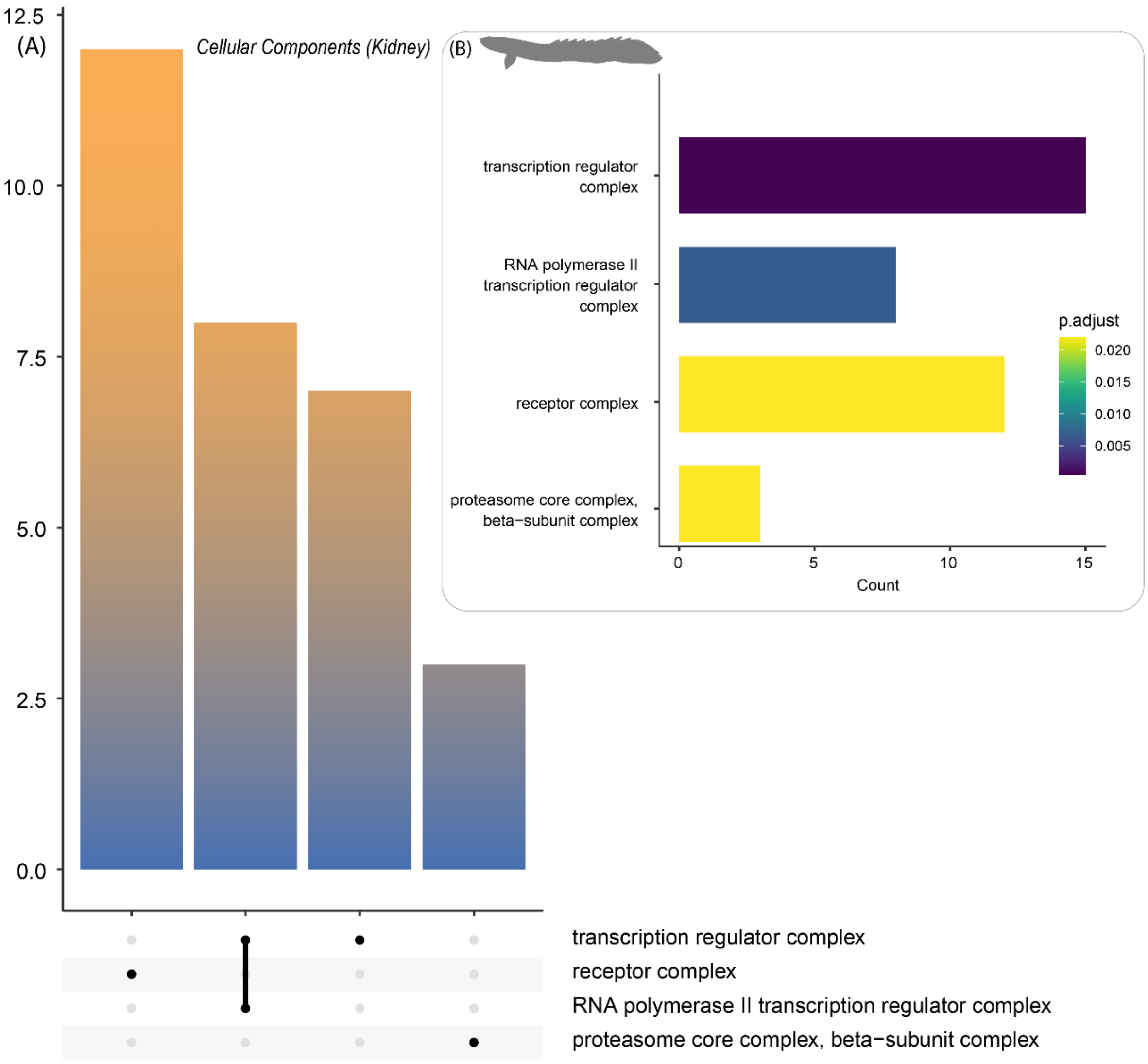


**Supplemental Figure S10:** Summary of Gene Ontology *Cellular Components* analysis from the *Polypterus bichir* kidney transcriptome.
Predicted proteins from the transcriptome were used as inputs to assess (A) processes and their intersections and (B) most common terms.

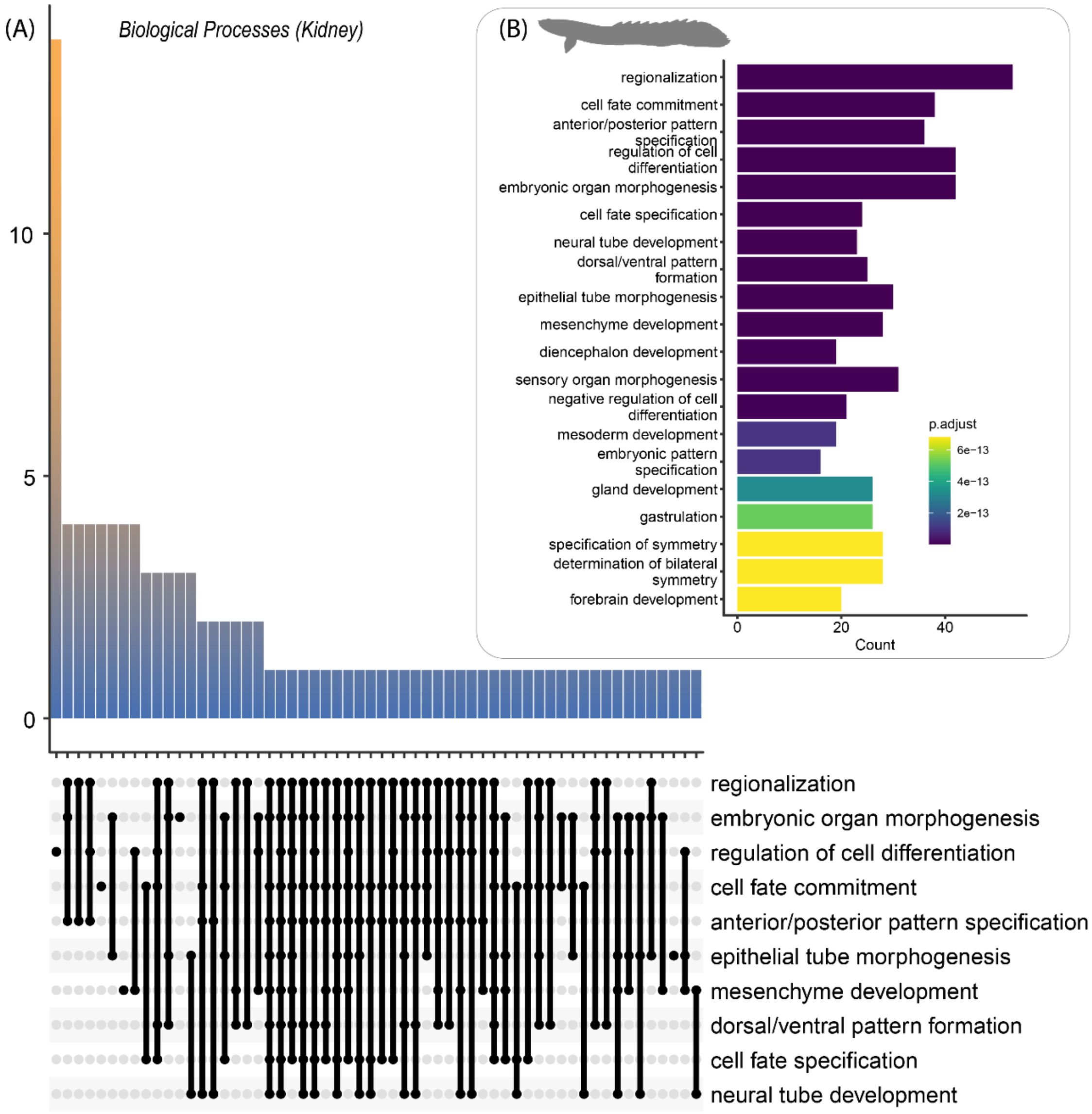


**Supplemental Figure S11:** Summary of Gene Ontology *Biological Processes* analysis from the *Polypterus bichir* kidney transcriptome.
Predicted proteins from the transcriptome were used as inputs to assess (A) processes and their intersections and (B) most common terms.

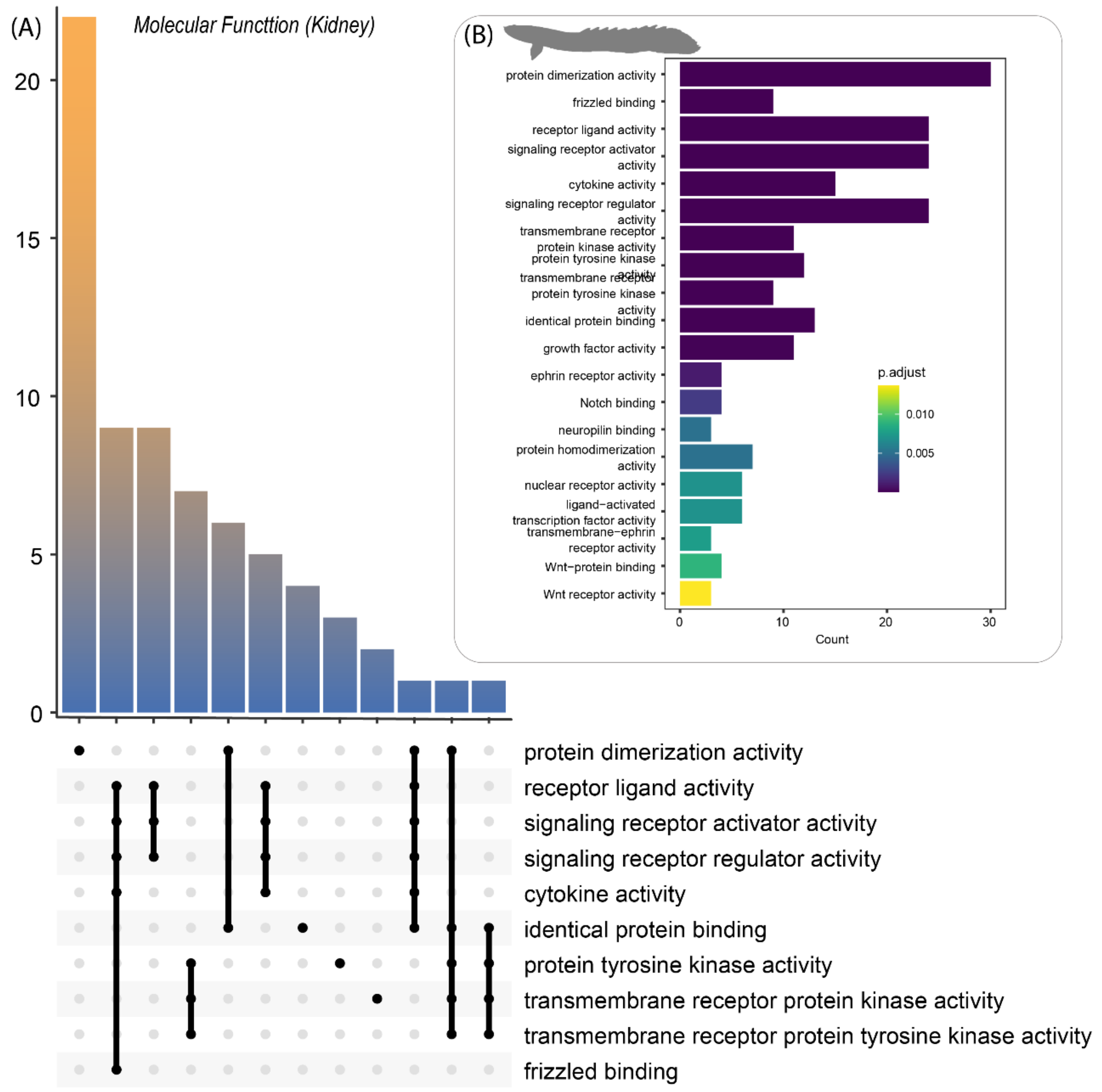


**Supplemental Figure S12:** Summary of Gene Ontology *Molecular Functions* analysis from the *Polypterus bichir* kidney transcriptome.
Predicted proteins from the transcriptome were used as inputs to assess (A) processes and their intersections and (B) most common terms.

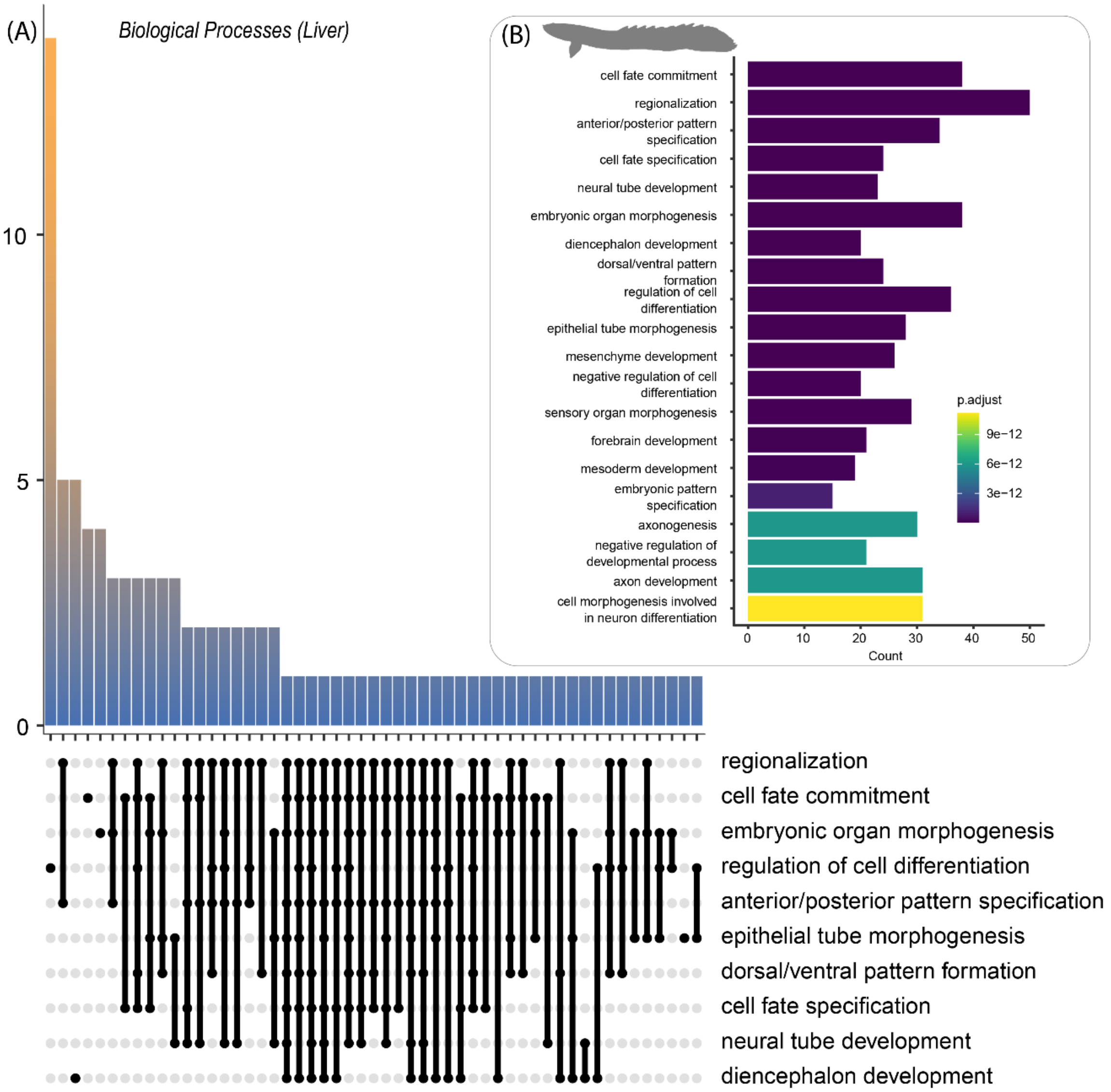


**Supplemental Figure S13:** Summary of Gene Ontology *Biological Processes* analysis from the *Polypterus bichir* liver transcriptome.
Predicted proteins from the transcriptome were used as inputs to assess (A) processes and their intersections and (B) most common terms.

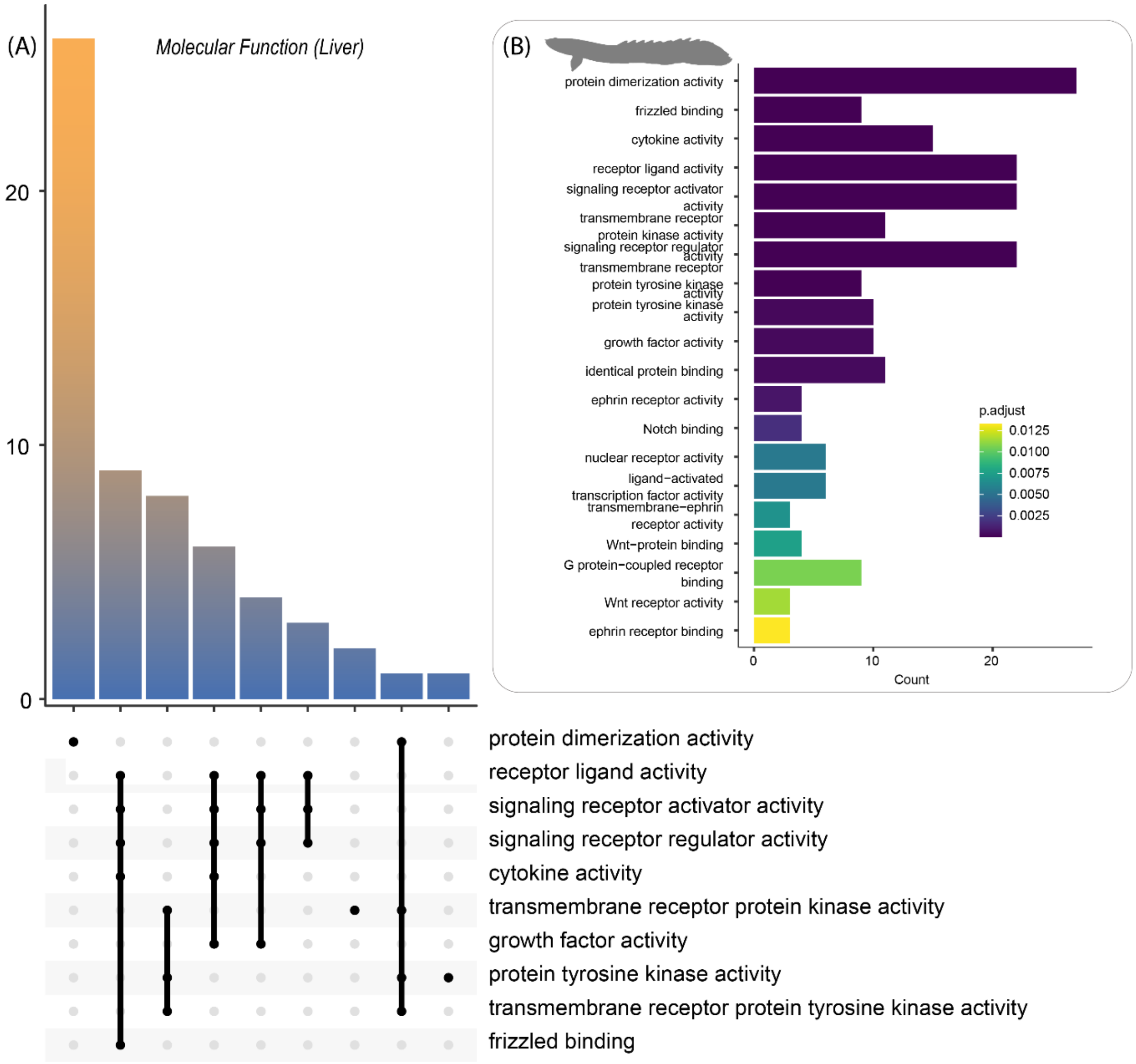


**Supplemental Figure S14:** Summary of Gene Ontology *Molecular Functions* analysis from the *Polypterus bichir* liver transcriptome.
Predicted proteins from the transcriptome were used as inputs to assess (A) processes and their intersections and (B) most common terms

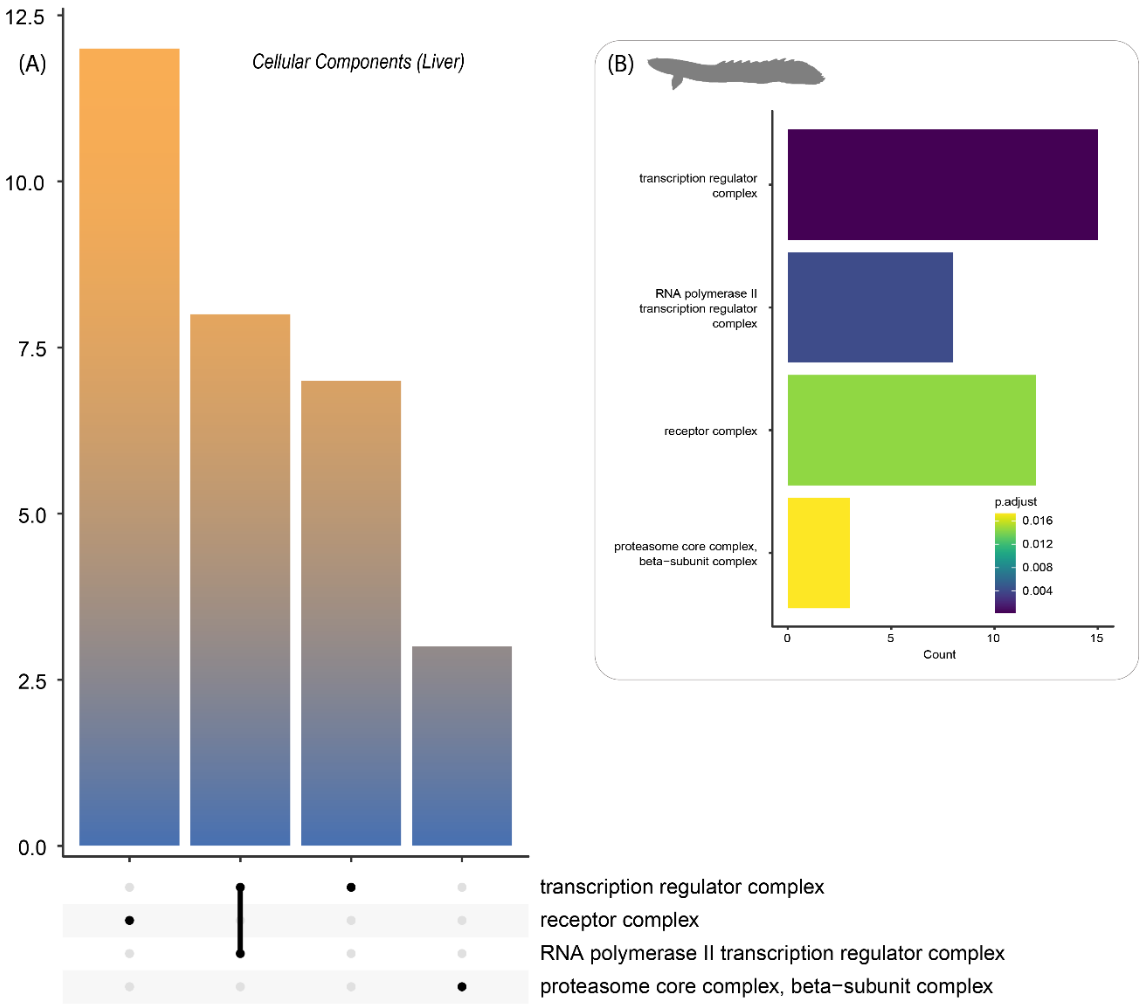


**Supplemental Figure S15:** Summary of Gene Ontology *Cellular Components* analysis from the *Polypterus bichir* liver transcriptome.
Predicted proteins from the transcriptome were used as inputs to assess (A) processes and their intersections and (B) most common terms.

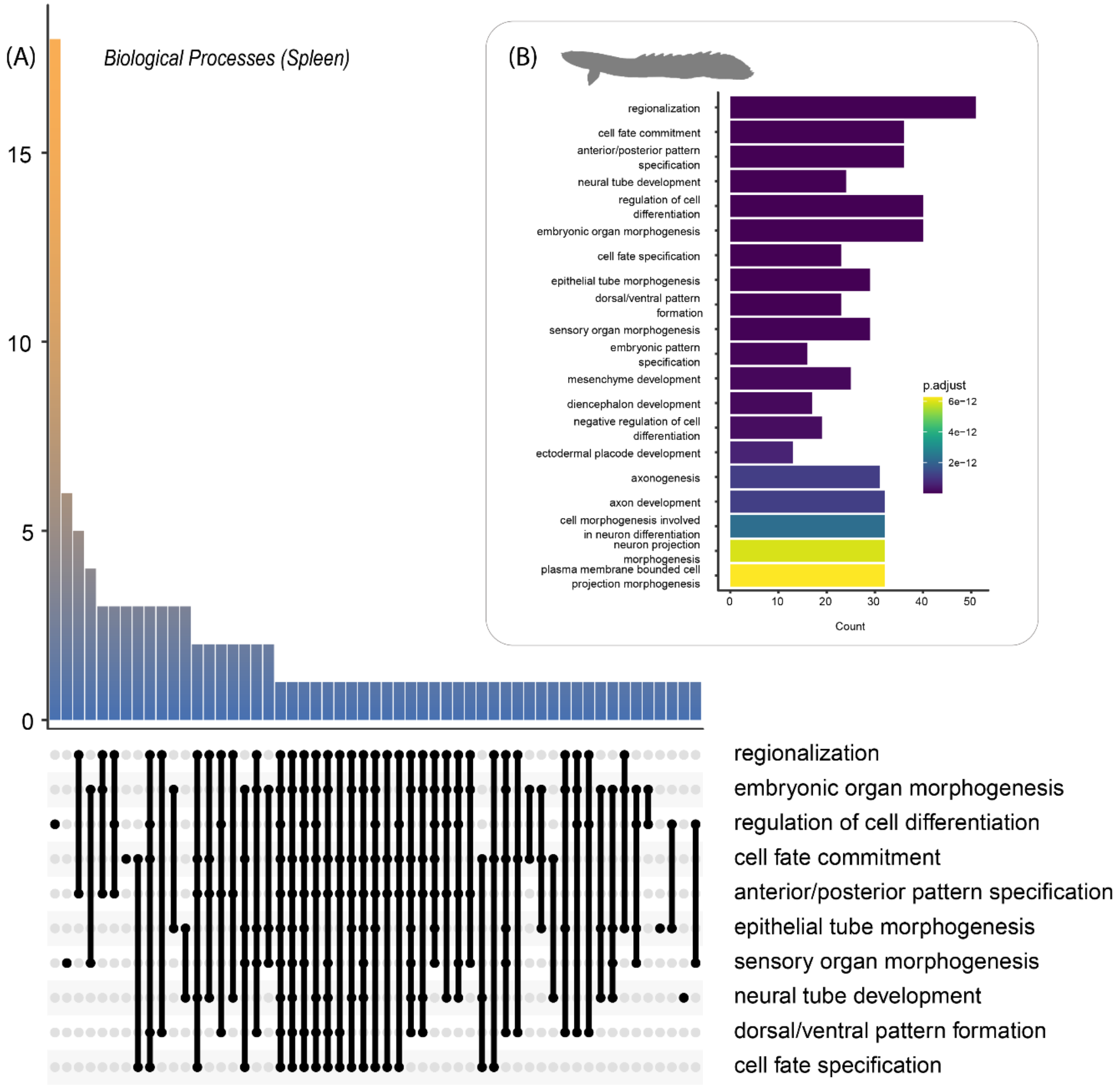


**Supplemental Figure S16:** Summary of Gene Ontology *Biological Processes* analysis from the *Polypterus bichir* spleen transcriptome.
Predicted proteins from the transcriptome were used as inputs to assess (A) processes and their intersections and (B) most common terms.

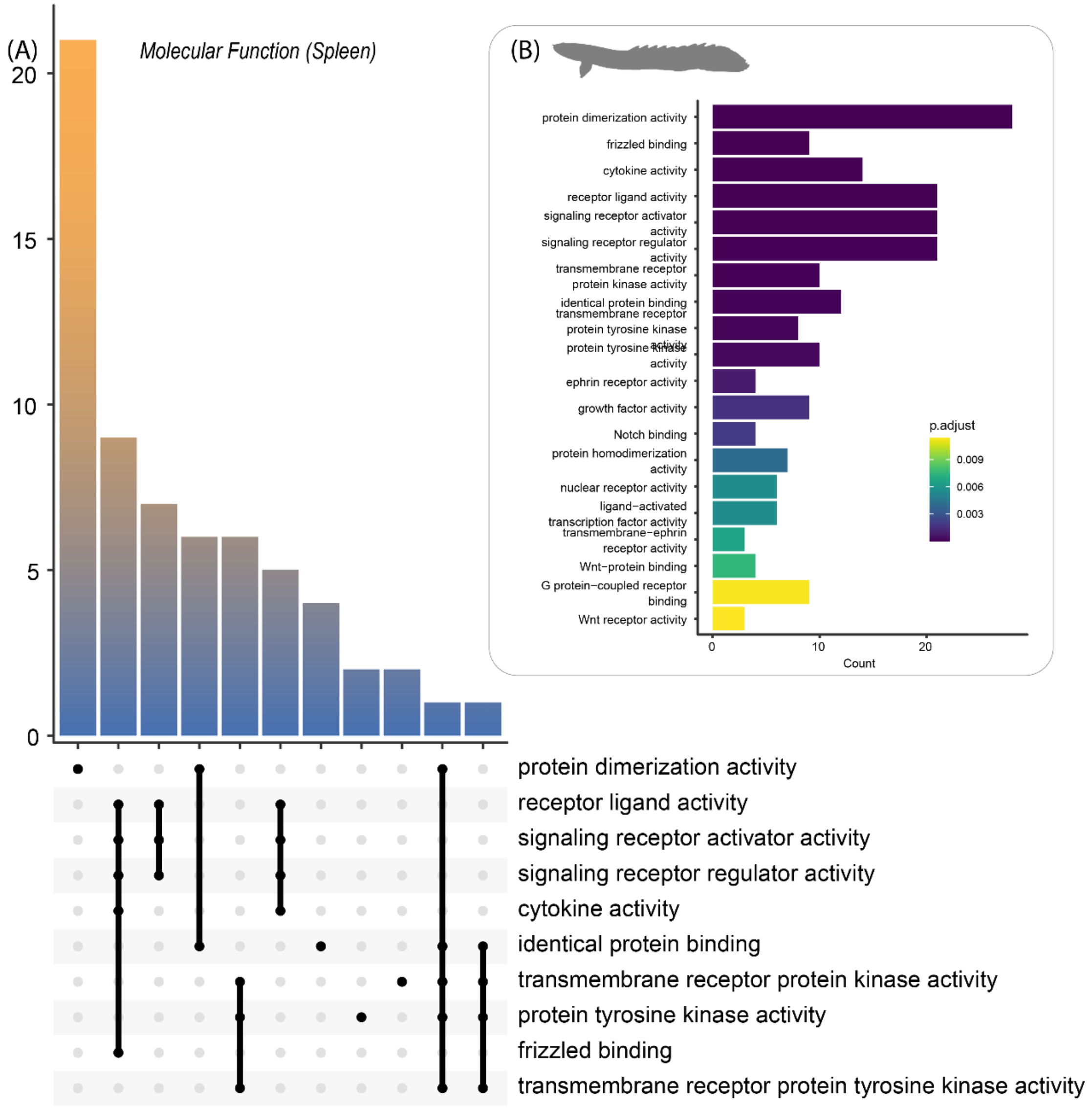


**Supplemental Figure S17:** Summary of Gene Ontology *Molecular Function* analysis from the *Polypterus bichir* spleen transcriptome.
Predicted proteins from the transcriptome were used as inputs to assess (A) processes and their intersections and (B) most common terms.

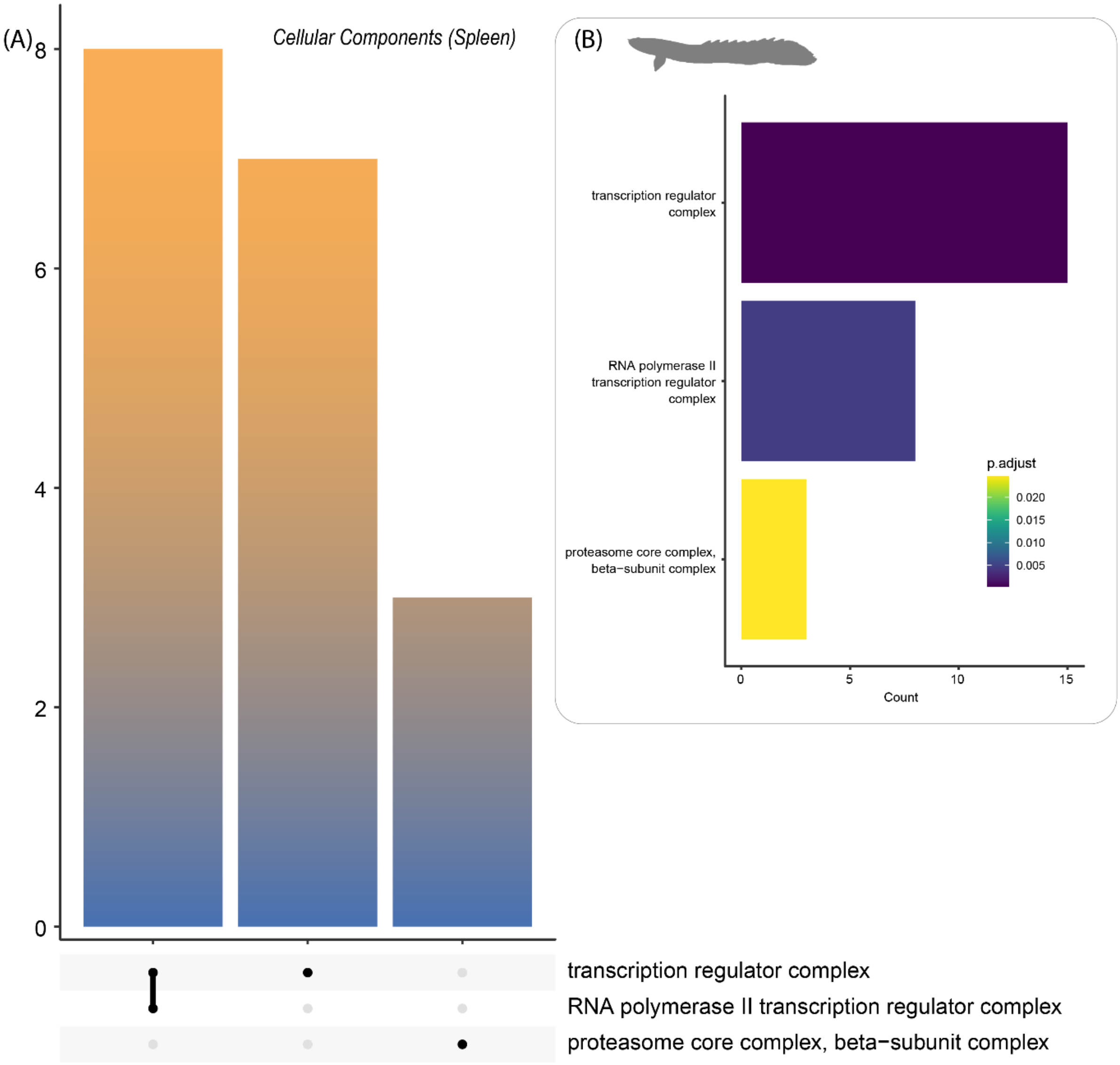


**Supplemental Figure S18:** Summary of Gene Ontology *Cellular Components* analysis from the *Polypterus bichir* spleen transcriptome.
Predicted proteins from the transcriptome were used as inputs to assess (A) processes and their intersections and (B) most common terms

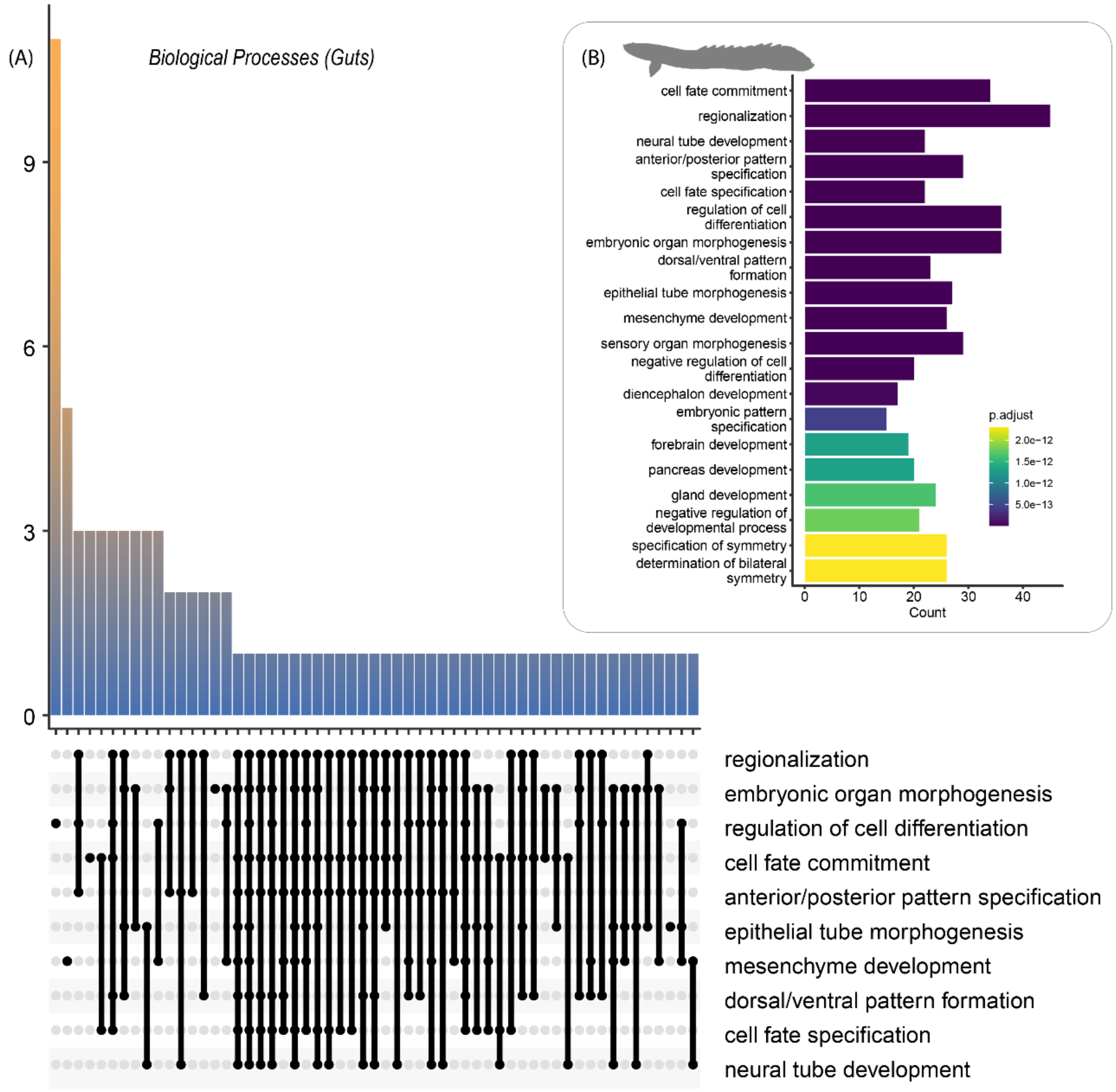


**Supplemental Figure S19:** Summary of Gene Ontology *Biological Processes* analysis from the *Polypterus bichir* gut transcriptome.
Predicted proteins from the transcriptome were used as inputs to assess (A) processes and their intersections and (B) most common terms.

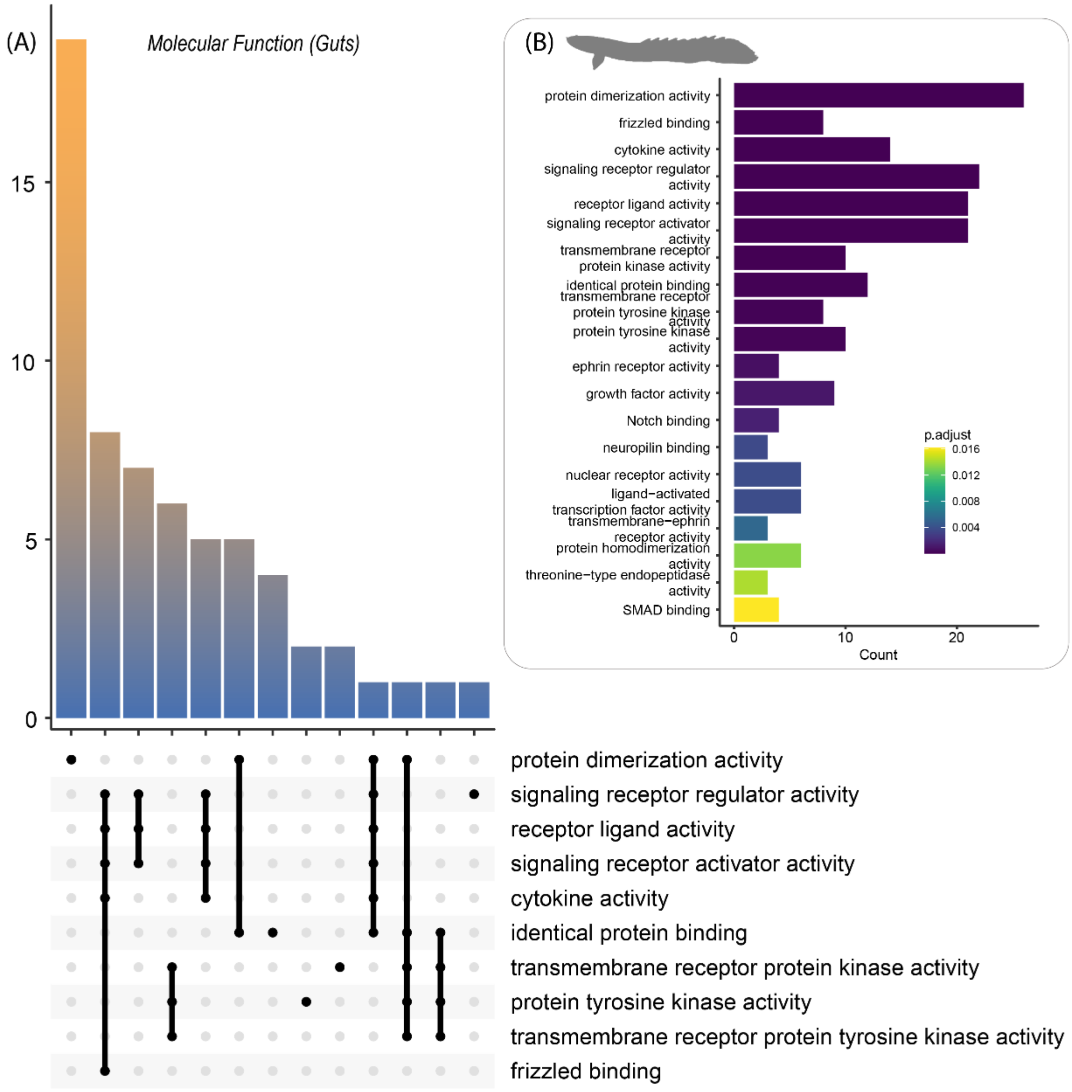


**Supplemental Figure S20:** Summary of Gene Ontology *Molecular Functions* analysis from the *Polypterus bichir* gut transcriptome.
Predicted proteins from the transcriptome were used as inputs to assess (A) processes and their intersections and (B) most common terms

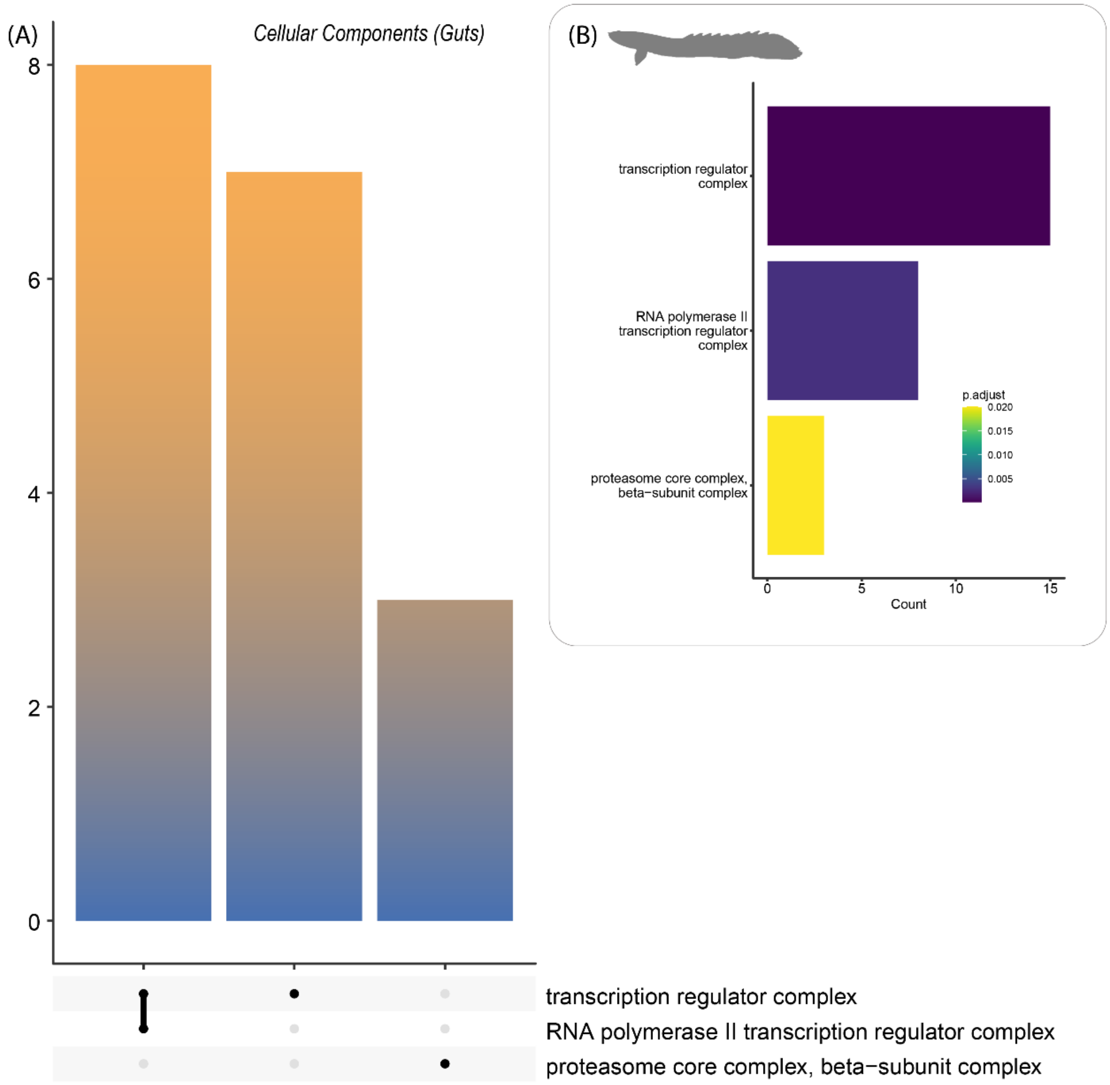


**Supplemental Figure S21:** Summary of Gene Ontology *Cellular Components* analysis from the *Polypterus bichir* gut transcriptome.
Predicted proteins from the transcriptome were used as inputs to assess (A) processes and their intersections and (B) most common terms.
.

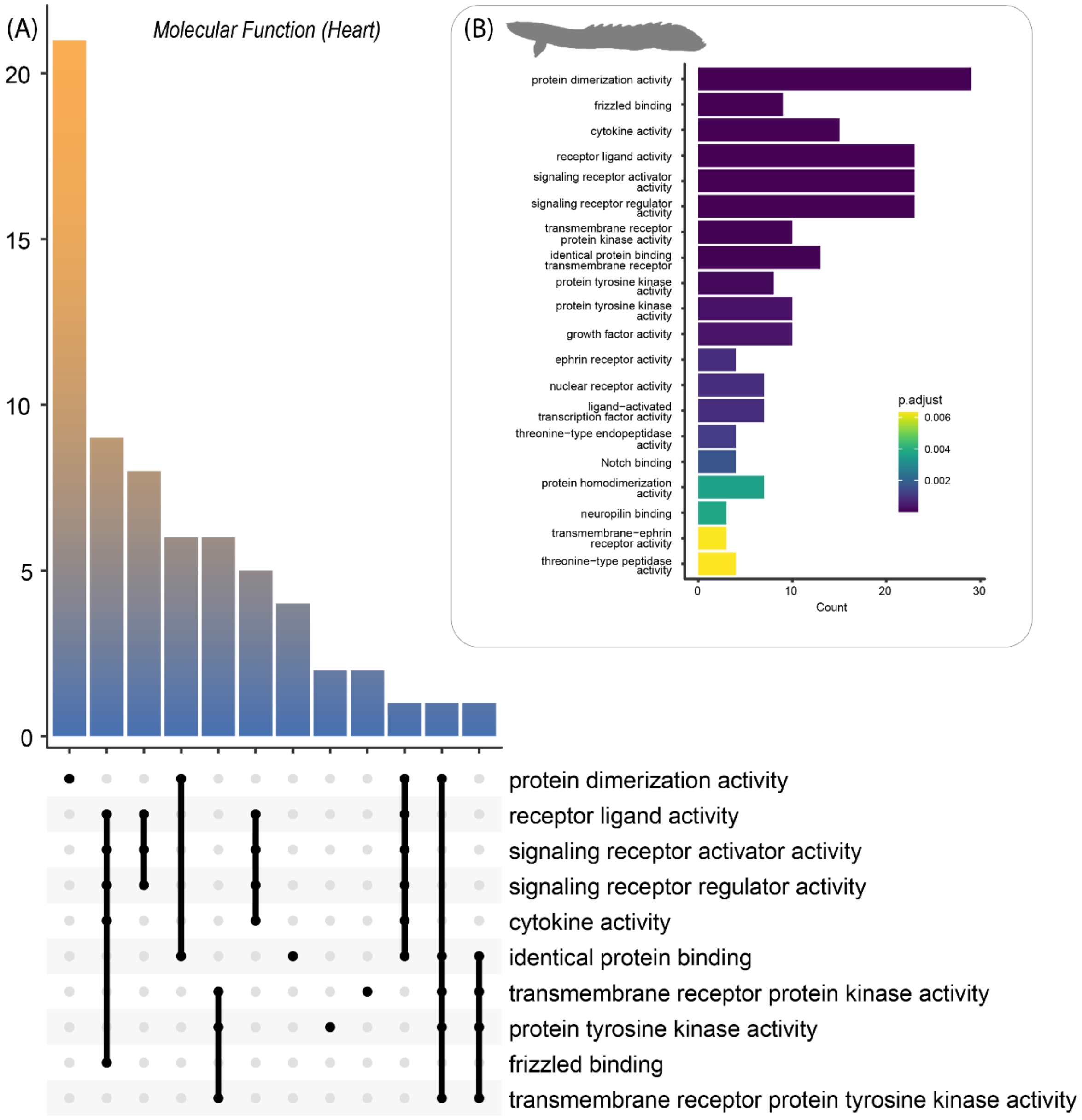


---

**Supplemental Figure S22:** Summary of Gene Ontology *Molecular Functions* analysis from the *Polypterus bichir* heart transcriptome.
Predicted proteins from the transcriptome were used as inputs to assess (A) processes and their intersections and (B) most common terms.

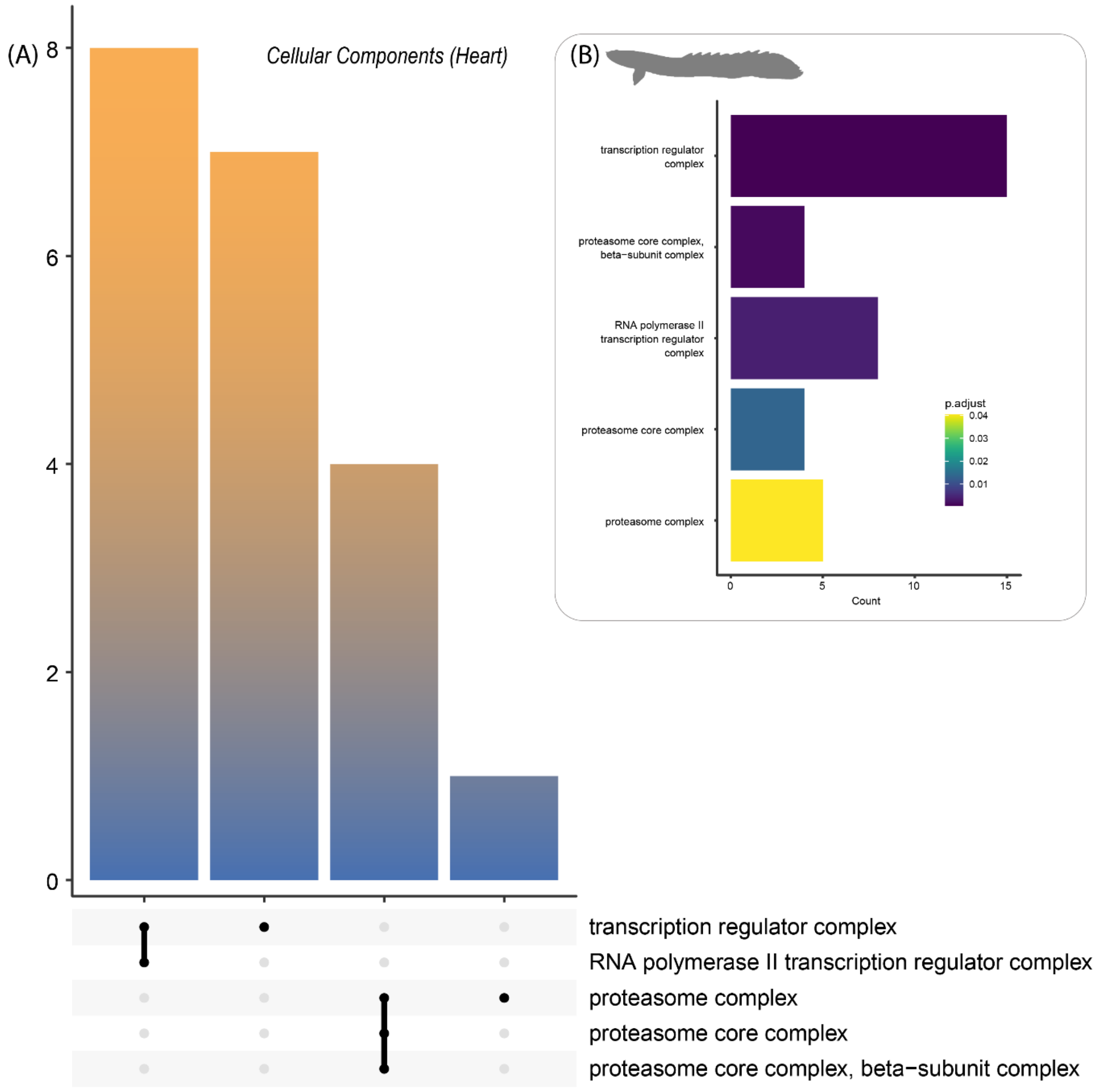


**Supplemental Figure S23:** Summary of Gene Ontology *Cellular Components* analysis from the *Polypterus bichir* heart transcriptome.
Predicted proteins from the transcriptome were used as inputs to assess (A) processes and their intersections and (B) most common terms.

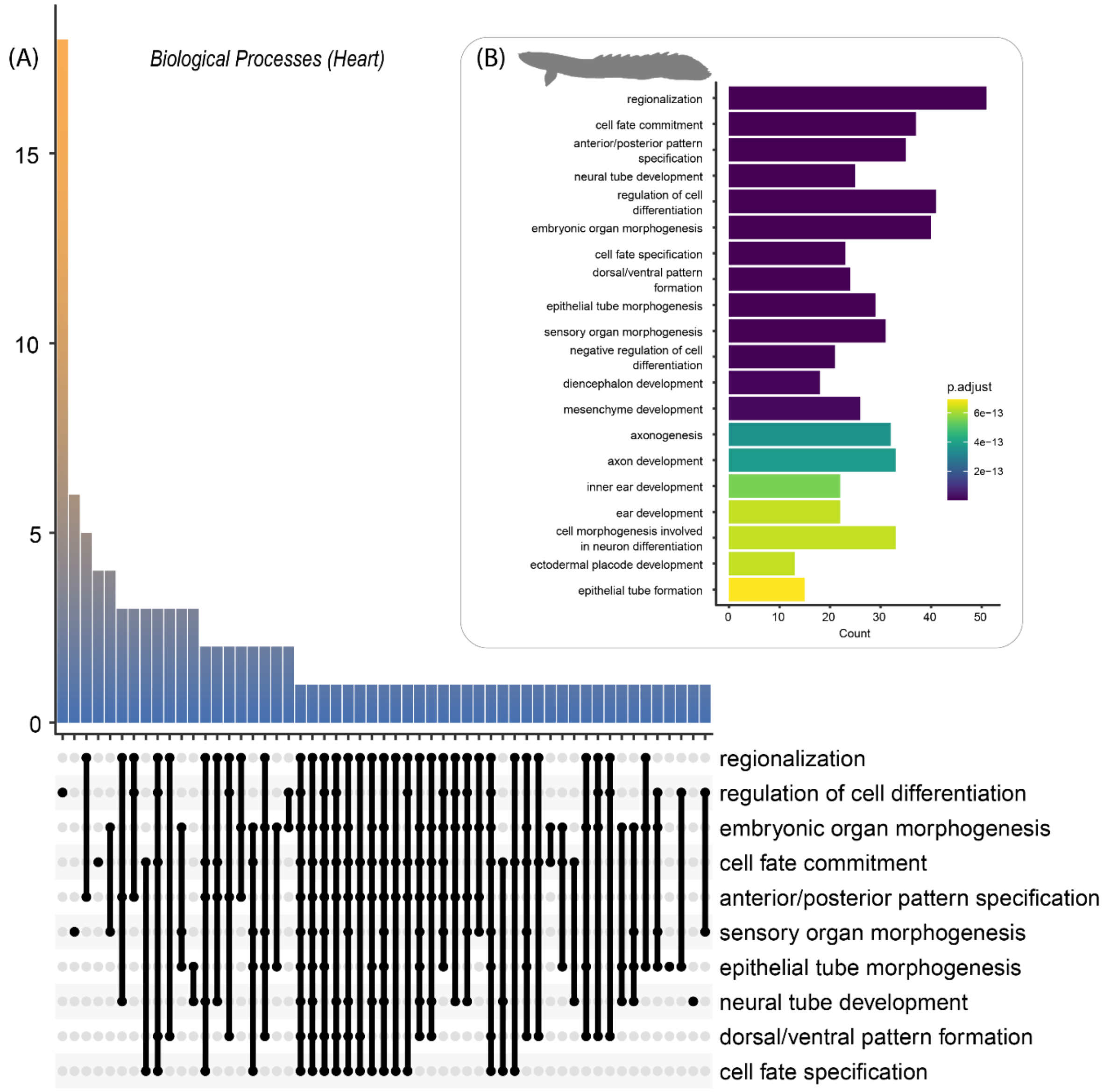


**Supplemental Figure S24:** Summary of Gene Ontology *Biological Processes* analysis from the *Polypterus bichir* heart transcriptome.
Predicted proteins from the transcriptome were used as inputs to assess (A) processes and their intersections and (B) most common terms.

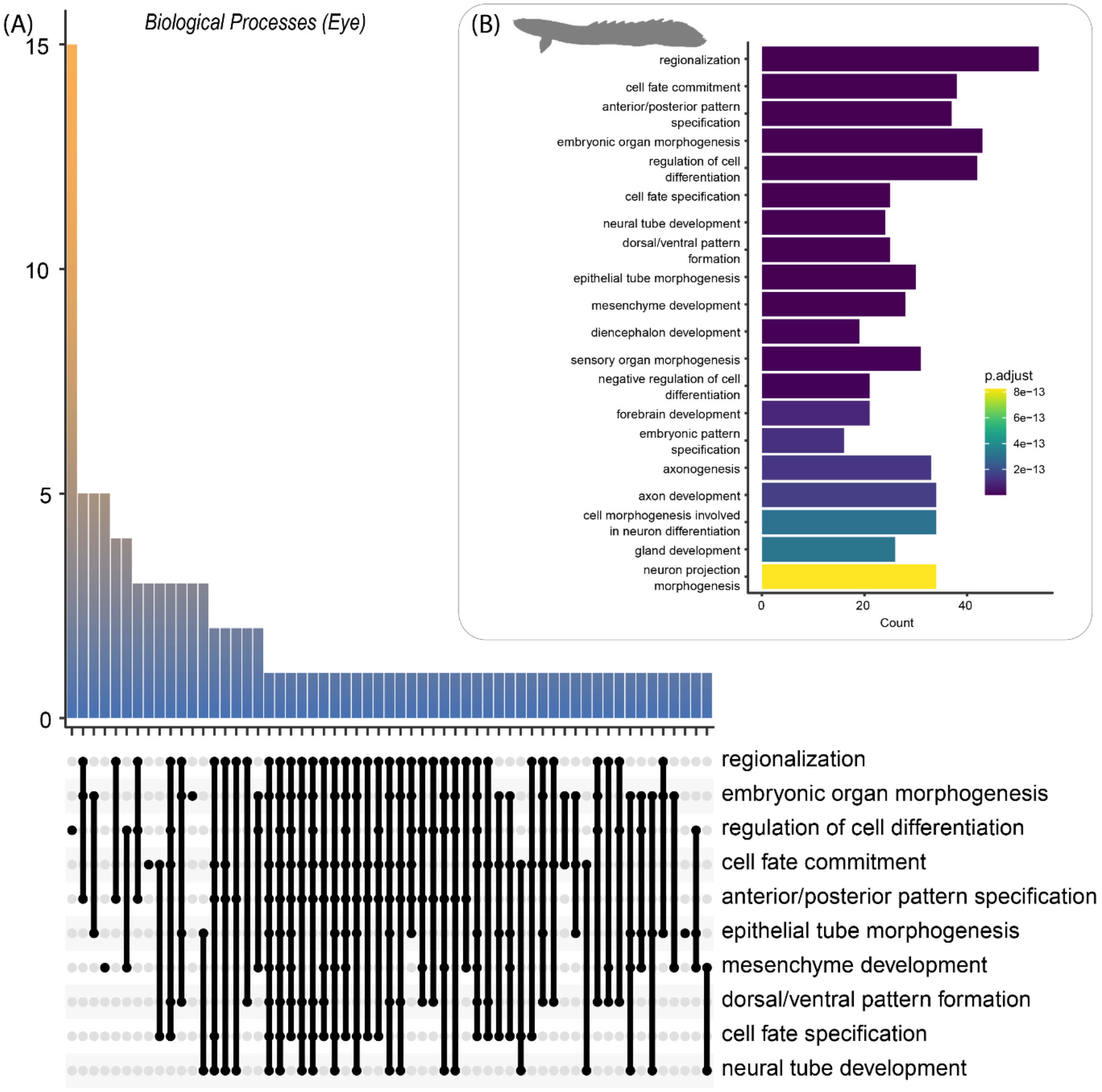


**Supplemental Figure S25:** Summary of Gene Ontology *Biological Processes* analysis from the *Polypterus bichir* eye transcriptome.
Predicted proteins from the transcriptome were used as inputs to assess (A) processes and their intersections and (B) most common terms.

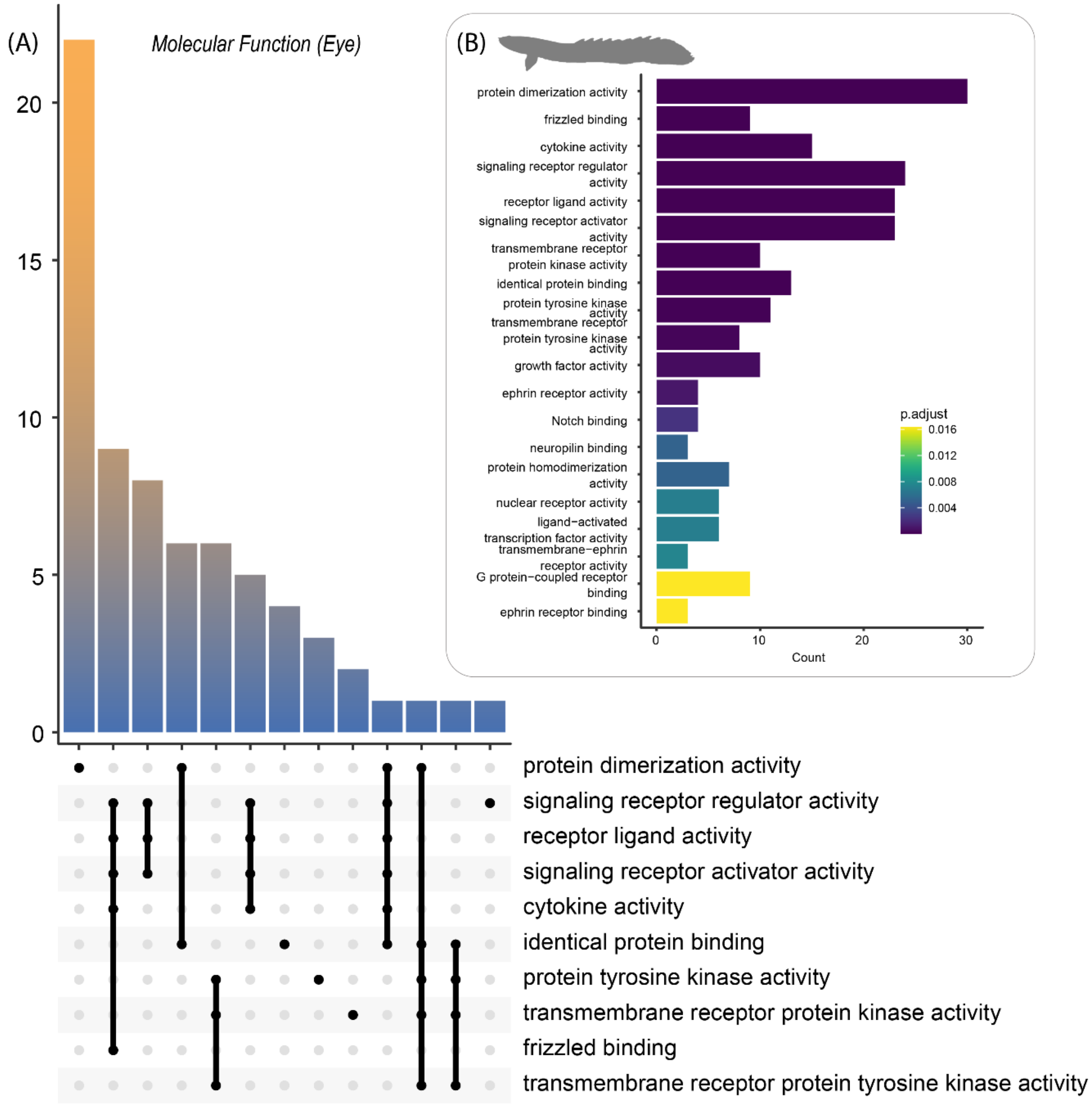


**Supplemental Figure S26:** Summary of Gene Ontology *Molecular Function* analysis from the *Polypterus bichir* eye transcriptome.
Predicted proteins from the transcriptome were used as inputs to assess (A) processes and their intersections and (B) most common terms.

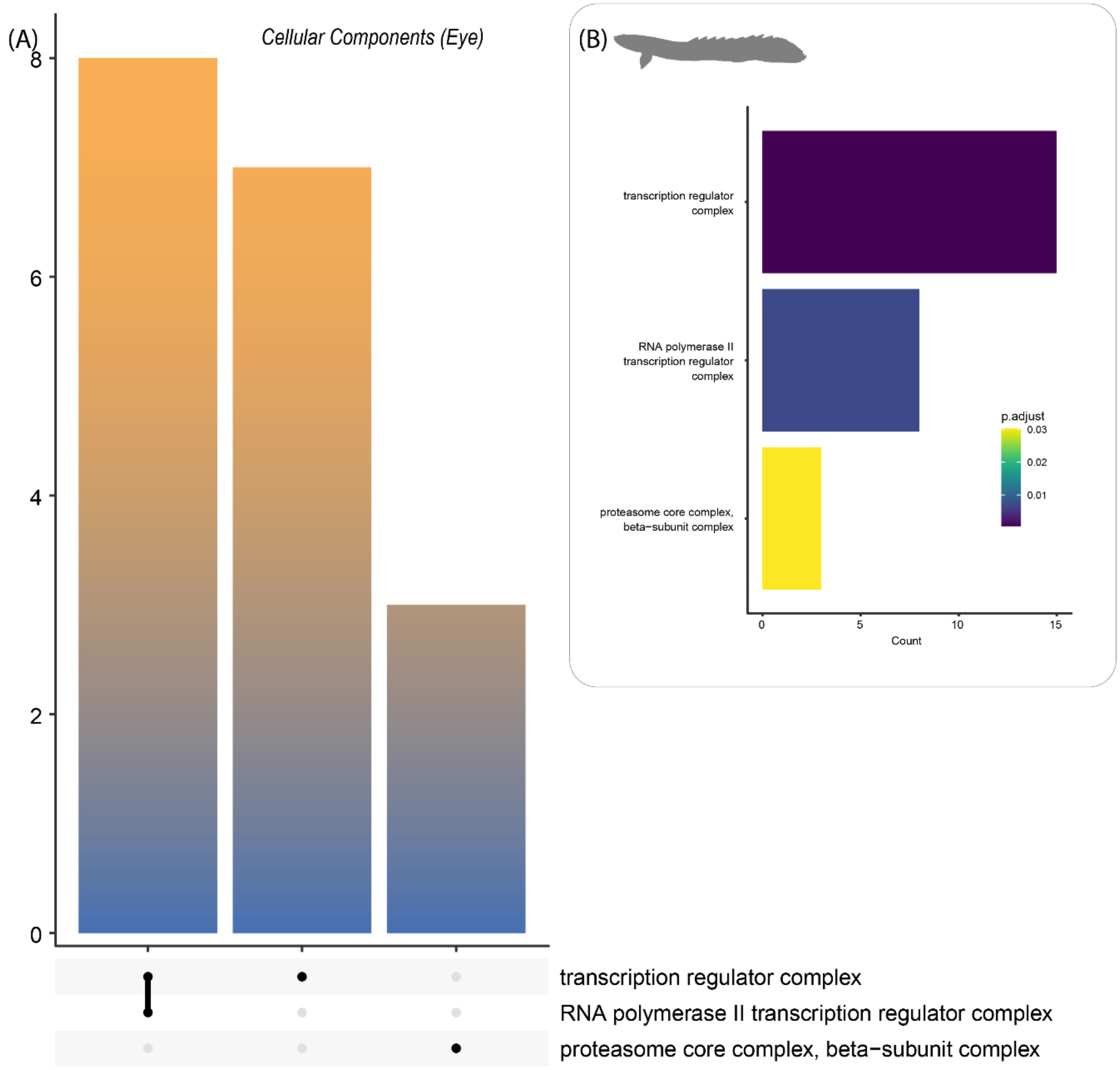


**Supplemental Figure S27:** Summary of Gene Ontology *Cellular Components* analysis from the *Polypterus bichir* eye transcriptome.
Predicted proteins from the transcriptome were used as inputs to assess (A) processes and their intersections and (B) most common terms.

**Supplemental Table S1.** Comparative genome assembly metrics of *Polypterus bichir* and related species

| Species | Genome size | Scaffold N50 | Scaffold L50 | Reference |
|---|---|---|---|---|
| *Polypterus bichir* | 3.9Gb | 202Mb | 7 | (This study) |
| *Erpetoichthys calabaricus* | 3.8Gb | 199Mb | 7 | (Ocampo Daza et al. 2021) |
| *Polypterus senegalus* | 3.6Gb | 189Mb | 8 | (Bi et al. 2021) |
| *Amia Calva* | 831Mb | 41Mb | 9 | (Thompson et al. 2021) |
| *Lepisosteus oculatus* | 945Mb | 6.9Mb | 45 | (Braasch et al. 2016) |
| *Lepisosteus osseus* | 930Mb | 53Mb | 8 | (Mallik et al. 2023) |

**Supplemental Table S2.** BUSCO scores of *Polypterus bichir*

| | Actinopterygii | Vertebrata |
|---|---|---|
| Complete BUSCOs (C) | 2630 (72.2%) | 2638 (78.77%) |
| Complete and single-copy BUSCOs (S) | 2563 (70.4%) | 2596 (77.4%) |
| Complete and duplicated BUSCOs (D) | 67 (1.8%) | 42 (1.3%) |
| Fragmented BUSCOs (F) | 90 (2.5%) | 272 (8.1%) |
| Missing BUSCOs (M) | 920 (25.3%) | 444 (13.2%) |
| Total BUSCO groups searched (n) | 3640 | 3354 |

**Supplemental Table S3.** Phylogenetic Signal and Variance Analysis using Pagel's Lambda, Blomberg's K, and ANOVA

| | **Pagel's lambda** | | **Blomberg's K** | | **ANOVA** | |
|---|---|---|---|---|---|---|
| | λ | *P*-value | K | *P*-value | F | *P*-value |
| DNA | 0.98 | **1.89e-13** | 0.48 | **6e-04** | 1.34 | 0.78 |
| LTR | 0.82 | **6.64e-11** | 0.41 | **1e-04** | 19 | 0.23 |
| LINE | 0.98 | **1.51e-19** | 0.94 | **1e-04** | 27 | 0.14 |
| SINE | 1.01 | **4.37e-15** | 0.53 | **3e-04** | 19 | 0.23 |
| Genome Size | 0.72 | **4.45e-15** | 0.66 | **6e-04** | **47.7** | **0.04** |

Bolded values indicate *P*-values below 0.05.

**Supplemental Table S4.** Evaluation of model fit for different TE types, genome size, and habitat characteristics under Brownian motion and Ornstein-Uhlenbeck (OU) models

| | **Brownian Motion** | | | **OU** | | |
|---|---|---|---|---|---|---|
| *Traits* | *Likelihood* | *AIC* | *AICw* | *Likelihood* | *AIC* | *AICw* |
| DNA | -276.21 | 556.43 | 0.12 | **-273.25** | **552.49** | **0.877** |
| LTR | -181.09 | 366.18 | <0.001 | **-165.23** | **336.47** | **>0.999** |
| LINE | ***-219.31*** | ***442.61*** | ***0.57*** | *-218.61* | *443.22* | *0.43* |
| SINE | ***-69.81*** | ***143.63*** | ***0.73*** | *-69.80* | *145.60* | *0.27* |
| Total TE | ***-360.08*** | ***724.17*** | ***0.61*** | *-359.53* | *725.07* | *0.39* |
| Size | *-601.31* | *1206.63* | *0.05* | ***-597.29*** | ***1200.58*** | ***0.95*** |
| Depth | *-528.20* | *1060.40* | *0.02* | ***-523.05*** | ***1052.10*** | ***0.98*** |
| Mean Latitude | *-481.17* | *966.34* | <0.001 | ***-466.67*** | ***939.35*** | ***>0.999*** |
| Median Latitude | *-498.38* | *1000.76* | <0.001 | ***-479.89*** | ***965.79*** | ***>0.999*** |
| Lower Latitude | *-505.02* | *1014.04* | <0.001 | ***-490.84*** | ***987.68*** | ***>0.999*** |
| Upper Latitude | *-494.66* | *993.32* | <0.001 | ***-467.44*** | ***940.89*** | ***>0.999*** |
| Genome Size | *-2224.41* | *4452.83* | <0.001 | ***-2208.42*** | ***4422.85*** | ***>0.999*** |

Bolded rows indicate best-fit model based on the highest AIC weight (AICw).

**Supplemental Table S5.** AIC scores of phylogenetic linear models evaluating each type of transposable element (TE)

| TE (y) Name | TE(x) Name | TE (x) | TE(x)+ Habitat | TE(x)+ Taxonomy | TE(x) + Taxonomy + Habitat |
|---|---|---|---|---|---|
| DNA | LTR | 534.53 | 537.62 | 536.22 | 539.19 |
| DNA | LINE | 534.30 | 537.20 | 535.54 | 538.46 |
| DNA | SINE | 551.10 | 552.68 | 552.85 | 560.85 |
| LTR | LINE | 309.25 | 307.19 | 311.22 | 309.19 |
| LTR | SINE | 368.15 | 366.66 | 369.91 | 366.36 |
| LINE | SINE | 440.59 | 443.64 | 441.93 | 445.01 |

**Supplemental Table S6.** Phylogenetic linear models depicting the correlation between TE abundance and genome size, latitude, body size, or depth under a Brownian motion model of trait evolution.

| **Trait** | TE | Intercept (a) | Coefficient (b) | SE (b) | T Value | *P*-value |
|---|---|---|---|---|---|---|
| Genome size | All | 32.72 | 2.65E-03 | 1.35E-03 | 1.97 | 0.05 |
| | DNA | 8.35 | 9.59E-10 | 6.00E-10 | 1.60 | 0.11 |
| | LINE | 5.96 | 4.80E-04 | 3.50E-04 | 1.39 | 0.17 |
| | LTR | 2.67 | 1.56E-10 | 2.41E-10 | 0.65 | 0.52 |
| | **SINE** | **0.62** | **1.89E-10** | **8.00E-05** | **2.37** | **0.02** |
| Latitude | All | 36.44 | 1.41E-02 | 3.92E-02 | 0.36 | 0.07 |
| | DNA | 9.50 | 1.20E-02 | 1.73E-02 | 0.69 | 0.49 |
| | LINE | 6.46 | 8.60E-03 | 9.96E-03 | 0.86 | 0.39 |
| | LTR | 2.60 | 1.12E-02 | 6.81E-03 | 1.64 | 0.10 |
| | **SINE** | **1.06** | **-6.06E-03** | **2.36E-03** | **-2.57** | **0.01** |
| Body Size | All | 36.15 | 1.55E-01 | 8.23E-01 | 0.19 | 0.85 |
| | DNA | 12.91 | 1.29E+01 | 3.58E-01 | -1.95 | 0.05 |
| | LINE | 5.56 | 2.60E-01 | 2.08E-01 | 1.25 | 0.22 |
| | LTR | 2.24 | 1.54E-01 | 1.44E-01 | 1.07 | 0.29 |
| | SINE | 0.77 | 3.16E-02 | 4.91E-02 | 0.64 | 0.52 |
| Depth | All | 30.63 | 6.62E-04 | 8.91E-04 | 0.74 | 0.46 |
| | DNA | 9.06 | 4.44E-04 | 2.88E-04 | 1.54 | 0.13 |
| | LINE | 4.04 | 5.10E-05 | 2.24E-04 | 0.22 | 0.82 |
| | LTR | 2.10 | -6.60E-06 | 1.09E-04 | -0.06 | 0.95 |
| | SINE | 0.67 | -5.50E-06 | 6.50E-05 | -0.08 | 0.93 |

Bolded rows indicate *P*-values below 0.05.

**Supplemental Table S7.** Results of phylogenetic linear models evaluating the correlation between TE abundance and genome size, latitude, body size, or depth under an OU model.

| Trait | TE | Intercept (a) | Coefficient (b) | SE (b) | T Value | P Value |
|---|---|---|---|---|---|---|
| Genome size | All | 20.22 | 1.15e-08 | 1.79e-09 | 6.42 | **4.55e-09** |
| | DNA | 4.76 | 4.15e-09 | 7.44e-10 | 5.58 | **2.05e-07** |
| | LINE | 1.45 | 3.21e-09 | 4.24e-10 | 7.58 | **1.75e-11** |
| | LTR | 0.09 | 1.01e-09 | 2.23e-10 | 4.55 | **1.52e-05** |
| | SINE | 0.04 | 2.51e-10 | 1.14e-10 | 2.21 | **2.95e-02** |
| Latitude | All | 1.52e-05 | -5.62e-03 | 0.06 | -0.09 | 0.93 |
| | DNA | 7.89 | 1.24e-03 | 0.02 | 0.05 | 0.96 |
| | LINE | 4.66 | -2.73e-02 | 0.01 | -1.79 | 0.07 |
| | LTR | 1.67 | 1.60e-04 | 0.01 | 0.02 | 0.98 |
| | SINE | 0.77 | -6.45e-03 | 0.01 | -1.93 | 0.06 |
| Body Size | All | 29.69 | -0.19 | 1.03 | -0.19 | 0.85 |
| | DNA | 11.18 | -0.89 | 0.40 | -2.20 | **0.03** |
| | LINE | 2.04 | 0.51 | 0.25 | 2.01 | **0.05** |
| | LTR | 0.59 | 0.29 | 0.11 | 2.56 | **0.01** |
| | SINE | 0.32 | 0.07 | 0.06 | 1.29 | 0.19 |
| Depth | All | 27.89 | 1.813e-03 | 1.13e-03 | 1.61 | 0.11 |
| | DNA | 7.24 | 1.50e-04 | 3.72e-04 | 0.40 | 0.69 |
| | LINE | 3.45 | 2.26e-04 | 3.14e-04 | 0.72 | 0.47 |
| | LTR | 1.58 | 4.3e-05 | 1.52e-04 | 0.28 | 0.78 |
| | SINE | 0.57 | 2.38e-05 | 8.11e-05 | 0.29 | 0.77 |

Bolded values indicate *P*-values below 0.05.

**Supplemental Table S8**. Evidence for correlated evolution between TEs from phylogenetic linear models

| **TE (x)** | TE (y) | Intercept (a) | Coefficient (b) | SE (b) | T Value | *P*-value | $R^2$ | $R^2$ Adj. |
|---|---|---|---|---|---|---|---|---|
| LINE | DNA | 8.07 | 1.94 | 0.71 | 2.73 | **7.53e-3** | 0.07 | 0.06 |
| LTR | DNA | 6.41 | 1.23 | 0.24 | 5.09 | **1.65e-06** | 0.20 | 0.20 |
| SINE | DNA | 4.51 | 0.80 | 0.15 | 5.18 | **1.16e-06** | 0.21 | 0.20 |
| LINE | LTR | 1.64 | 0.54 | 0.22 | 2.46 | **0.01549** | 0.06 | 0.05 |
| SINE | LTR | -0.14 | 0.46 | 0.05 | 8.83 | **3.36e-14** | 0.44 | 0.43 |
| SINE | LINE | 4.70 | 1.12 | 0.40 | 2.78 | **0.006573** | 0.07 | 0.06 |

Bolded values indicate *P*-values below 0.05.

**References:**

Bi X, Wang K, Yang L, Pan H, Jiang Haifeng, Wei Q, Fang M, Yu H, Zhu C, Cai Y, et al. 2021. Tracing the genetic footprints of vertebrate landing in non-teleost ray-finned fishes. *Cell* 184:1377-1391.e14.

Braasch I, Gehrke AR, Smith JJ, Kawasaki K, Manousaki T, Pasquier J, Amores A, Desvignes T, Batzel P, Catchen J, et al. 2016. The spotted gar genome illuminates vertebrate evolution and facilitates human-teleost comparisons. *Nat. Genet.* 48:427–437.

Mallik R, Carlson KB, Wcisel DJ, Fisk M, Yoder JA, Dornburg A. 2023. A chromosome-level genome assembly of longnose gar, Lepisosteus osseus. *G3* [Internet] 13. Available from: http://dx.doi.org/10.1093/g3journal/jkad095

Ocampo Daza D, Bergqvist CA, Larhammar D. 2021. The Evolution of Oxytocin and Vasotocin Receptor Genes in Jawed Vertebrates: A Clear Case for Gene Duplications Through Ancestral Whole-Genome Duplications. *Front. Endocrinol.* 12:792644.

Thompson AW, Hawkins MB, Parey E, Wcisel DJ, Ota T, Kawasaki K, Funk E, Losilla M, Fitch OE, Pan Q, et al. 2021. The bowfin genome illuminates the developmental evolution of ray-finned fishes. *Nat. Genet.* 53:1373–1384.
